# SENP1 in the retrosplenial agranular cortex regulates core autistic-like symptoms in mice

**DOI:** 10.1101/2021.01.24.427868

**Authors:** Kan Yang, Yuhan Shi, Xiujuan Du, Jincheng Wang, Yuefang Zhang, Shifang Shan, Yiting Yuan, Ruoqing Wang, Chenhuan Zhou, Yuting Liu, Zilin Cai, Yanzhi Wang, Liu Fan, Huatai Xu, Juehua Yu, Jinke Cheng, Fei Li, Zilong Qiu

## Abstract

Autism spectrum disorder (ASD) is a highly heritable neurodevelopmental disorder, in which core symptoms are defects of social interaction and evidently repetitive behaviors. Although around 50-70 % of ASD patients have comorbidity of intellectual disabilities (ID) or developmental delay (DD), there are some ASD patients who exhibit only core symptoms but without ID/DD, raising the question whether there are genetic components and neural circuits specific for core symptoms of ASD. Here, by focusing on ASD patients who do not show compound ID or DD, we identified a *de novo* heterozygous gene-truncating mutation of the Sentrin-specific peptidase1 (*SENP1*) gene, coding the small ubiquitin-like modifiers (SUMO) deconjugating enzyme, as a potentially new candidate gene for ASD. We found that *Senp1* haploinsufficient mice exhibited core symptoms of autism such as deficits in social interaction and repetitive behaviors, but normal learning and memory ability. Moreover, we found that the inhibitory and excitatory synaptic functions were severely affected in the retrosplenial agranular (RSA) cortex of *Senp1* haploinsufficient mice. Lack of *Senp1* led to over SUMOylation and degradation of fragile X mental retardation protein (FMRP) proteins, which is coded by the *FMR1* gene, also implicated in syndromic ASD. Importantly, re-introducing SENP1 or FMRP specifically in RSA fully rescued the defects of synaptic functions and core autistic-like symptoms of *Senp1* haploinsufficient mice. Together, these results demonstrated that disruption of the SENP1-FMRP regulatory axis in the RSA may cause core autistic symptoms, which provide a candidate brain region of ASD for potential therapeutic intervene by neural modulation approaches.

## Introduction

Autism spectrum disorders (ASD) is a group of neurodevelopment disorders, characterized by defects in social communication and stereotypic behaviors (Lord et al., 2020). Although ASD is one of the highest heritable mental disorders (Sandin et al., 2017; Wang et al., 2017), the genetic landscape of ASD was not revealed until the application of genome-wide sequencing technologies such as whole-exome sequencing and whole-genome sequencing (De Rubeis et al., 2014; Iossifov et al., 2014). The major contributory genetic components of ASD include *de novo* variants and rare inherited variants, as well as chromosomal structural variations (Iakoucheva et al., 2019; Searles Quick et al., 2021). Among various genetic variants, the *de novo* variants of large effect, for example protein-truncating variants, would account for 10-15% of total contributing genetic causes of ASD and provide the first line of clue for illustrating the neural mechanisms responsible for disrupted social behaviors of ASD patients (Searles Quick et al., 2021).

In the recent ASD genetic study using the largest cohort so far, researchers proposed that there were a group of ASD candidate genes primarily affecting core symptoms of autism such as social communication and repetitive behavior (referred to as ASD predominant genes), whereas another group of ASD candidate genes also led to defects in cognition caused by neurodevelopmental delay (referred to as ASD & NDD genes) (Satterstrom et al., 2020). Although whether “ASD predominant” and “ASD & NDD” genes could be clearly separated is still arguable (Myers et al., 2020), focusing on the ASD candidate genes mainly affecting symptoms in social domains, instead of cognitive functions, may provide novel insights for understanding the genomics and neural mechanisms underlying social behaviors in mammals.

In a whole-exome sequencing study on over 600 ASD trios in Shanghai Xinhua Hospitals, we identified a *de novo* heterozygous protein-truncating mutation in the *SENP1* gene of an autism patient with impaired social functions, but without developmental delay (normal Intelligence Quotient (IQ)/ Developmental Quotient (DQ)). SENP1 (Sentrin-specific peptidase1) plays a pivotal role in post-translational SUMOylation modifications by releasing SUMO groups from proteins (Flotho and Melchior, 2013). Homozygous mutations of the *SENP1* gene have been implicated in severe neurometabolic diseases (Tarailo-Graovac et al., 2016). Although SENP1 plays critical roles in regulating various physiological functions including metabolism, as well as neural injury (Sun et al., 2020b; Wang et al., 2019), the role for SENP1 in the central nervous system remains largely unclear (Choi et al., 2016; Sun et al., 2014).

In this study, we showed that *Senp1* haploinsufficient mice (*Senp1*^+/-^) exhibited defects of social behaviors and increased stereotypic behaviors, without deficiency in learning and memory tasks. Interestingly, we found that the inhibitory neurons were specifically affected in the retrosplenial agranular (RSA) cortex of *Senp1* ^+/-^ mice. The inhibitory and excitatory synaptic transmission of layer II/III pyramidal neurons in the RSA region were altered in the *Senp1* haploinsufficient mice comparing to wild-type littermates. The orchestrating alteration of inhibitory and excitatory synaptic transmission was also seen in *Cntnap3* knockout mice, another autism mouse model reported by our lab (Tong et al., 2019), suggesting that the co-regulation of inhibitory and excitatory synapse may be one of the key signatures of autism pathophysiology. Mechanistically, we found that lack of SENP1 led to the over SUMOylation and degradation of fragile X syndrome mental retardation protein (FMRP) in RSA. Remarkably, we could rescue the defects of the social behaviors of Senp1 haploinsufficiency mice by introducing SENP1 or FMRP specifically into the RSA region. These data indicate that *SENP1* is a candidate gene for “ASD predominant gene” affecting primarily social functions and the RSA region may serve as a circuitry node implicated in regulating mammalian social behaviors. Interestingly, it is recently reported that retrosplenial granular cortex (RSG) is also responsible for social behavior deficits of *Fmr1* knockout mice (Shang et al., 2021). Together, these findings support the notion that the retrosplenial cortex plays a critical role in regulating social behaviors of mice.

## Results

### Identification of a *de novo* mutation in the *SENP1* gene in a ASD proband and autistic-like behaviors in *Senp1* haploinsufficient mice

In order to study ASD candidate genes in Chinese cohorts, we performed whole-exome sequencing to over 600 ASD probands with their parents to search for *de novo* variants which may contribute to ASD symptoms. All ASD probands were further examined with Childhood Autism Rating Scale (CARS), Autism Diagnostic Observation Schedule (ADOS) and Intelligence Quotient (IQ)/ developmental Quotient (DQ) tests to acquire the comprehensive appearances of social and cognitive functions. We identified a *de novo* protein-truncating mutation in the *SENP1* gene (NM001267594:c.151C>T:p.Q51X), validated by Sanger sequencing (Figure 1A). The mutation takes place at a conserved region of the *SENP1* gene across mammals, which lead to premature termination of the SENP1 protein during translation (Figure 1B, C).

**Figure 1.**
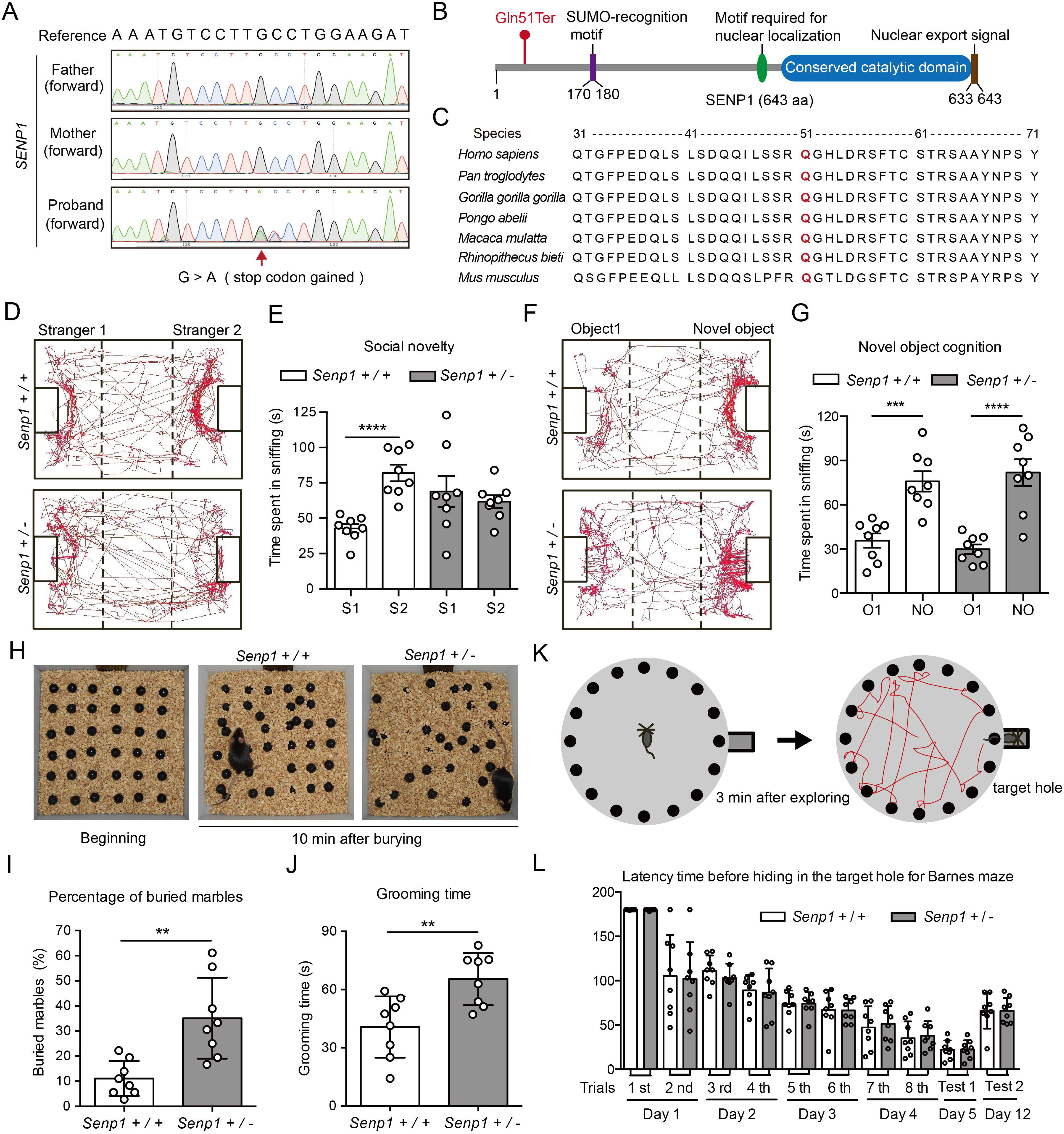
*Senp1*-haploinsufficient mice exhibit impaired social novelty behaviors and elevated repetitive restrictive behaviors. (A) A *de novo* stop-codon gained mutation in human *SENP1* gene verified by Sanger sequencing. Sanger sequencing traces of unaffected parents and affected proband were aligned to human reference sequence. (B) Schematic illustration of premature termination of human SENP1 protein. Red fonts showed the mutated site (Gln 51 to termination codon). (C) Conserved amino acid sequences of SENP1 protein across mammals. (D) Representative locomotor traces of mice in the three-chamber test for social novelty. (E) Quantification of time mice spent in sniffing familiar and novel partners (n = 8, for each genotype). (F) Representative locomotor traces of mice in the three-chamber novel object cognition. (G) Quantification of time mice spent in sniffing familiar and novel object (n = 8, for each genotype). (H) Representative images showing the beginning and the end of the marble burying test. (I) Quantification of percentage of buried marbles within 10 min (n = 8, for each genotype). (J) Quantification of grooming time within 30 min (n = 8, for each genotype). (K) Schematic illustration for a mouse to perform a trial within 3 min of Barnes maze test. (L) Quantification of latency time before hiding in the target hole (n = 8, for each genotype). ** *p* < 0.01, *** *p* <0.001, **** *p* < 0.0001. n is the number of mice used in each experiment. Bars represent means ± SD. See also Figure S1 and Figure S2.

The male patient carrying the SENP1 mutation exhibited typical deficits in social behaviors including communication and interaction, measured by ADOS and CARS scores (Table S1). Interestingly, the patient exhibited largely normal Developmental Quotient measured by the Gesell development scale (Table S1), suggesting that mutation of *SENP1* specifically affected the behaviors in social domain, instead of cognition domain, in human.

To investigate roles of the *SENP1* gene in social behavior, we use laboratory mouse as an animal model to examine whether genetic deletion of the *Senp1* gene may affect social behaviors in mice (Wang et al., 2019). Since homozygous deletion of the *Senp1* gene causes lethality of mouse, we examine a battery of behavioral tests in the *Senp1* haploinsufficient mice which carry heterozygous deletion of the *Senp1* gene (Figure S1A). We first examined the anxiety level using the open field test and found that *Senp1* haploinsufficient mice exhibited the same level of anxiety as wild-type (WT) littermates (Figures S1B, S2A-S2E). We next examine the social behaviors of *Senp1* haploinsufficient mice along with WT littermates in the classic three-chamber test (Figure S1C). We found that although *Senp1* haploinsufficient mice showed the same social approach feature comparing to WT littermates (Figure S2F-S2I), *Senp1* haploinsufficient mice displayed significant defects in the social novelty test, by not showing increased preference with novel partners in comparison with WT littermates (Figure 1D, E). This suggests that lack of *Senp1* may lead to abnormal social behaviors. To further validate whether *Senp1* haploinsufficient mice are able to recognize novel objects, we used the object cognition test (Figure S1D) and found no significant difference in object cognition preference between *Senp1* haploinsufficient and WT mice (Figure S2J-S2M). Similarly, no difference in the novel object cognition test was found (Figure 1F, G). This data suggests that *Senp1* haploinsufficient mice have specific defects in social cognition behaviors.

Stereotypic and repetitive behaviors are also core symptoms of ASD. Thus, we examined whether *Senp1* haploinsufficient mice have stereotypic and repetitive behaviors via the restrictive and repetitive behaviors test (Figure S1E). Interestingly, *Senp1* haploinsufficient mice showed more stereotypic behaviors by measuring the marbles buried and times for self-grooming (Figure 1H-J). Lastly, we investigated the spatial learning ability of mutant and WT mice by using the Barnes maze test (Figure S1F). We found that *Senp1* haploinsufficient mice exhibited the normal spatial learning ability as WT mice (Figure 1K, 1L; Figure S2N). Together, these results indicate that *Senp1* haploinsufficient mice display abnormal social behaviors but normal object cognition and spatial learning ability, which is in consistent with clinical symptoms of the ASD patient carrying *SENP1* mutant and suggests that SENP1 in the brain may specifically contribute to the social interaction behaviors.

### Compromised brain development in *Senp1* haploinsufficient mice

To examine whether *Senp1* haploinsufficiency may lead to abnormal neural development, we first analyzed whether brain structures were altered due to loss of *Senp1*. Surprisingly, we found that the brain width and thickness of cerebral cortex significantly increased in both male and female *Senp1* haploinsufficient mice at 2 months age, comparing to WT littermates (Figure 2A-2D). Brain overgrowth is an intriguing symptom often seen in ASD patients and ASD mouse models such as PTEN mutant mice (Kwon et al., 2006).

**Figure 2.**
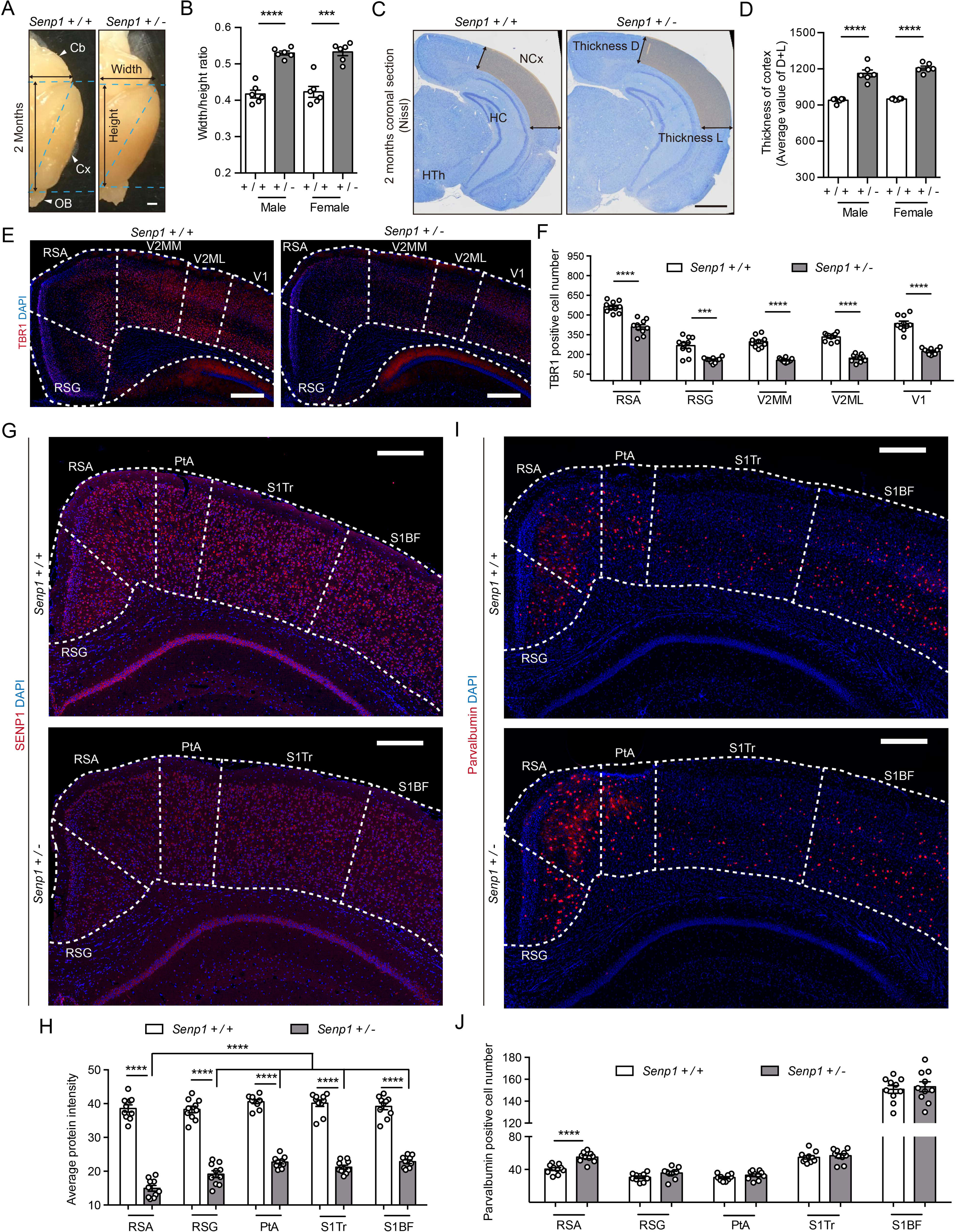
*Senp1*-haploinsufficient mice show abnormal development of brain and RSA-specific increase of parvalbumin positive interneuron. (A) Representative pictures of whole-mount brains for each genotype at 2 months of age. Cb, cerebellum; Cx, cortex; OB, olfactory bulb. Scale bar, 1mm. (B) Quantification of the width/height ratio for brains of each gender (n = 6, for each genotype). (C) Representative coronal sections of brain visualized with Nissl staining from both genotypes. NCx, neocortex; HC, hippocampus; HTh, hypothalamus; Thickness D, thickness for dorsal edge of neocortex; Thickness L, thickness for lateral edge of neocortex. Scale bar, 1mm. (D) Quantification of average thickness of neocortex for each gender (n = 3, for each genotype). (E) Representative coronal immunofluorescence staining sections of brain stained for anti-TBR1 (red) and DAPI (blue) at 3 months. RSA, retrosplenial agranular cortex; RSG, retrosplenial granular cortex; V2MM, secondary visual cortex, mediomedial area; V2ML, secondary visual cortex, mediolateral area; V1, primary visual cortex. Scale bar, 400µm. (F) Quantification of TBR1 positive cell number within each cortical region (n = 3, for each genotype). (G) Representative coronal immunofluorescence staining sections of cortex stained for anti-SENP1 (red) and DAPI (blue) at 3 months of age. RSA, retrosplenial agranular cortex; RSG, retrosplenial granular cortex; PtA, parietal association cortex; S1Tr, primary somatosensory cortex, trunk region; S1BF, primary somatosensory cortex, barrel field. Scale bar, 400µm. (H) Quantification of average protein intensity within each cortical region (n = 1, for each genotype). (I) Representative coronal immunofluorescence staining sections of brain stained for parvalbumin (red) and DAPI (blue) at 3 months. RSA, retrosplenial agranular cortex; RSG, retrosplenial granular cortex; PtA, parietal association cortex; S1Tr, primary somatosensory cortex, trunk region; S1BF, primary somatosensory cortex, barrel field. Scale bar, 400µm. (J) Quantification of parvalbumin positive cell number within each region (n = 3, for each genotype). *** *p* < 0.001, **** *p* < 0.0001. n is the number of mice used in each experiment. Bars represent means ± SD.

To further investigate how the brain structure is altered in *Senp1* haploinsufficient mice, we examine expression levels of various neuronal markers in the brain. First, we found that the expression of TBR1, an important cortical marker gene and also a ASD risk gene, was dramatically decreased in *Senp1* haploinsufficient mice across various cortices at 2 months of age, in comparison with WT mice (Fazel Darbandi et al., 2018; Huang et al., 2014; Huang et al., 2019) (Figure 2E, 2F), suggesting that the cortical development program was impaired in the *Senp1* haploinsufficient mice.

We next determined the expression pattern of SENP1 in the mouse brain. Using immunostaining with antibody against SENP1, we found that the expression of SENP1 was highly enriched in various cortical regions of the WT mouse brain and decreased in the brain of *Senp1* haploinsufficient mice (Figure 2G, 2H). Notably, we found that the downregulation of SENP1 proteins in the *Senp1* haploinsufficient mice was mostly significant in the retrosplenial agranular (RSA) cortex, a subregion of retrosplenial cortex (RSC), comparing to other cortices (Figure 2G, 2H).

Interestingly, the numbers of two major types of GABAergic neurons, parvalbumin (PV) and somatostatin (SST) positive neurons, were specifically increased in the RSA cortex of *Senp1* haploinsufficient mice, comparing to WT mice (Figure 2I, 2J, Figure S3A-S3E). In particular, the number of PV and SST positive neurons was increased in the RSA layer II/III and layer V of the *Senp1* haploinsufficient mice, in comparison with WT mice (Figure S3A, S3B, S3F, S3G), suggesting that the inhibitory synaptic connections in RSA may be altered in *Senp1* haploinsufficient mice.

Next to posterior cingulate cortex, the retrosplenial cortex (RSC), composing of two parts — RSA (retrosplenial agranular) and RSG (retrosplenial granular), is implicated in the top-down modulation of sensorimotor information from primary sensory cortices (Bicanski and Burgess, 2020), as well as spatial navigation due to extensive connections to hippocampal formation (Vann et al., 2009). Interestingly, researchers recently reported that ketamine treatment in mouse specifically activated the slow wave oscillation in the retrosplenial cortex (Vesuna et al., 2020). Furthermore, the rhythmic activation of deep layer neurons (layer V) in the retrosplenial cortex causally led to a series of dissociation-like phenotype in mice, including withdraw of social interaction (Vesuna et al., 2020). Since defects in sensorimotor functions are prominent in ASD patients, it is plausible that the RSC may regulate social behavior via modulating sensorimotor information integration. We hypothesized that abnormal synaptic connections in the RSA region may cause deficits of social interaction behaviors, due to incapable of properly processing sensory information to higher centers.

### Disrupted inhibitory and excitatory synaptic functions in the RSA neurons of *Senp1* haploinsufficient mice

We next used immunostaining and electrophysiology to examine whether *Senp1* haploinsufficiency alters the structure and function of GABAergic synapses in the RSA cortex (Figure 3A-3D). To examine the GABAergic synapses formed on RSA neurons in WT and *Senp1* haploinsufficient mice, we injected two adeno-associated viruses (AAV-CAG-FLEX-EGFP, AAV-hSyn-Cre) to sparsely label layer II/III neurons in the RSA region and imaged the synapses formed on the soma of labelled neurons by con-focal microscopy (Figure 3A, 3B). We found that numbers of inhibitory synapses, measured by punctum co-labelled with vesicular GABA transporter (VGAT) and Gepyrin, were significantly increased in soma of RSA layer II/III neurons of *Senp1* haploinsufficient mice, comparing to WT mice (Figure 3E-3G). Consistently, the punctum co-labelled with VGAT and the α1 subunit of GABA_A_ receptors were also increased in RSA layer II/III neurons of *Senp1* haploinsufficient mice, in comparison with WT mice, suggesting that *Senp1* haploinsufficiency leads to more inhibitory synapses formation on RSA layer II/III neurons (Figure 3H-3J).

**Figure 3.**
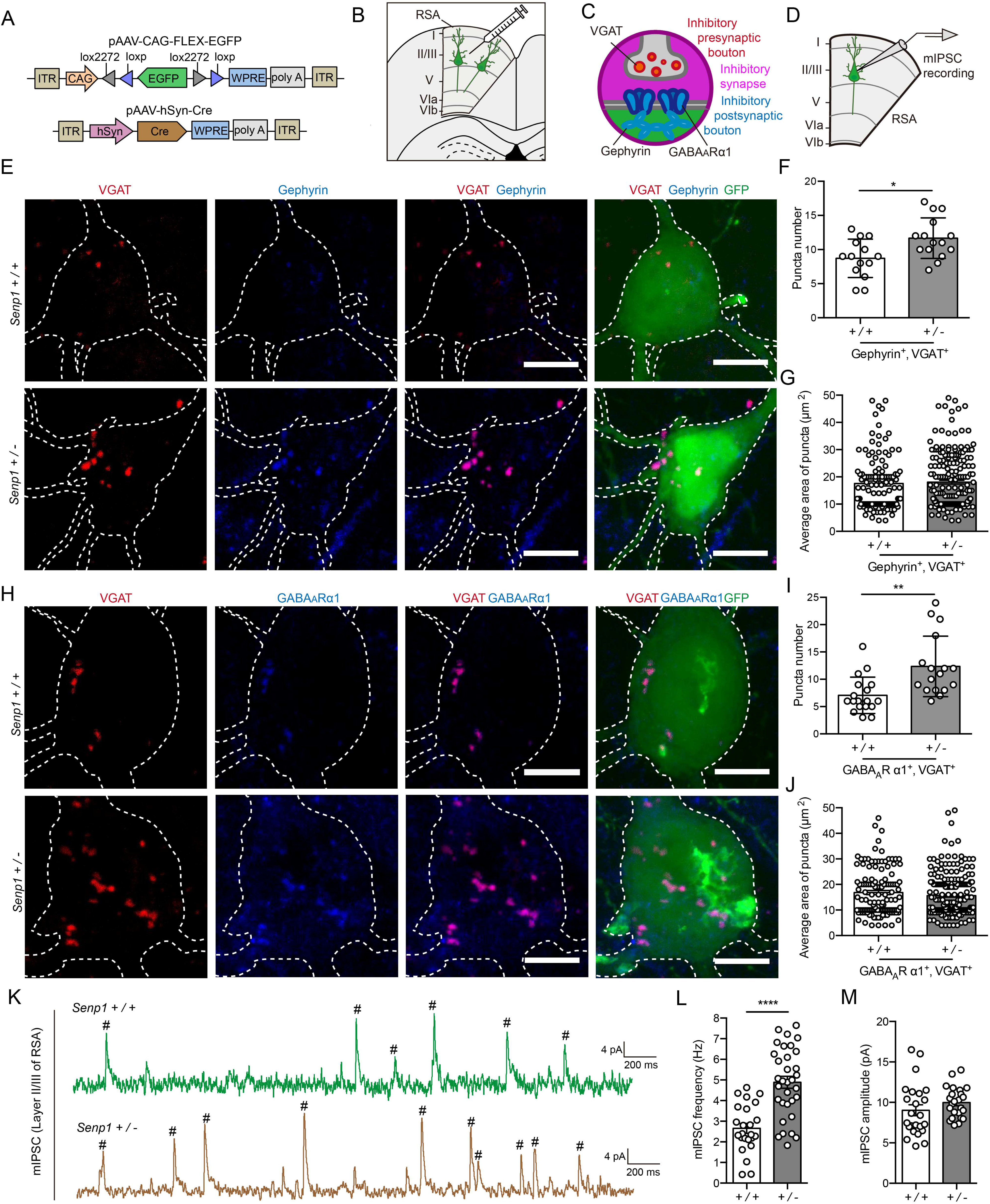
Increased of inhibitory synapses on pyramidal neuron correlates to elevated mIPSC frequency for *Senp1*-haploinsufficient mice. (A) Schematic illustration of AAV viral vector design strategy. (B) Schematic illustration for AAV injection to sparsely label pyramidal neurons in layer II/III of RSA. (C) Schematic illustration of immunofluorescence staining strategy to mark the inhibitory synapse, stained for inhibitory presynaptic marker, VGAT (red), and inhibitory postsynaptic marker, Gephyrin (blue), and inhibitory postsynaptic marker, GABA_A_Rα1 (blue). (D) Schematic illustration of electrophysiological recording on layer II/III pyramidal neurons of RSA brain slices. (E) Representative immunofluorescence staining pictures for inhibitory synapses on soma part of sparsely labeled pyramidal neuron in RSA, stained for VGAT (red), Gephyrin (blue), and GFP (green). Scale bar, 60µm. (F) Quantification of puncta number of both Gephyrin and VGAT positive bouton on the soma (neuron number n = 14, from 3 *Senp1 ^+/+^* mice; n = 15, from 3 *Senp1 ^+/-^* mice). (G) Quantification of average area for the puncta (≥ 4µm^2^) of both Gephyrin and VGAT positive bouton on the soma (punctum number n = 122, from 14 neurons calculated in Figure 3F, for *Senp1 ^+/+^* mice; n = 210, from 15 neurons calculated in Figure 3F, for *Senp1 ^+/-^* mice). (H) Representative immunofluorescence staining pictures for inhibitory synapses on soma part of sparsely labeled pyramidal neuron in RSA, stained for VGAT (red), GABA_A_Rα1 (blue), and GFP (green). Scale bar, 60µm. (I) Quantification of puncta number of both GABA_A_Rα1 and VGAT positive bouton on the soma (neuron number n = 18, from 3 *Senp1 ^+/+^* mice; n = 17, from 3 *Senp1 ^+/-^* mice). (J) Quantification of average area for the puncta (≥ 4µm^2^) of both GABA_A_Rα1 positive and VGAT positive bouton on the soma (punctum number n = 127, from 18 neurons calculated in Figure 3I, for *Senp1 ^+/+^* mice; n = 210, from 17 neurons calculated in Figure 3I, for *Senp1 ^+/-^* mice). (K) Representative mIPSC traces of pyramidal neurons in the layer II/III of RSA. (L) Quantification of mIPSC frequency for pyramidal neurons in the layer II/III of RSA (neuron number n = 27 from 3 *Senp1 ^+/+^* mice; n = 30 from 3 *Senp1 ^+/-^* mice). (M) Quantification of mIPSC amplitude for pyramidal neurons in the layer II/III of RSA (neuron number n = 27 from 3 *Senp1 ^+/+^* mice; n = 30 from 3 *Senp1 ^+/-^* mice). * *p* < 0.05, ** *p* < 0.01, **** *p* < 0.0001. Bars represent means ± SD for Figure 3F and 3I; bars represent means ± SEM for Figure 3G, 3J, 3L and 3M.

To further address whether the inhibitory synaptic transmission of RSA layer II/III neurons of *Senp1* haploinsufficient mice was altered, we performed the whole-cell patch clamp recording on the layer II/III pyramidal neurons in RSA brain slices from mice of both genotypes. We found that the frequency of miniature inhibitory post-synaptic currents (mIPSCs) in RSA layer II/III neurons of *Senp1* haploinsufficient mice was significantly higher than that in WT mice (Figure 3K-3M). Together, this data indicates that inhibitory synaptic transmission of RSA layer II/III pyramidal neurons in *Senp1* haploinsufficiency mice was increased likely due to more inhibitory synapses formed on these neurons.

We then wonder whether *Senp1* haploinsufficiency may also alter the structure and functions of excitatory synapses in RSA neurons. We performed sparse labeling by AAV injections followed by immunostaining experiments and electrophysiology (Figure 4A-4D). Interestingly, we found that the numbers of excitatory synapses, represented by punctum labeled by co-staining of PSD95 and vesicular glutamate transporter 1 (VGlut1), as well as co-staining of glutamate ionotropic receptor AMAP type subunit 2 (GRIA2) and VGlut1, were dramatically decreased in soma of RSA layer II/III neurons of *Senp1* haploinsufficient mice, comparing to WT mice (Figure 4E-4J). In consistent, we found that the frequency, but not amplitude, of miniature excitatory postsynaptic current (mEPSCs) significantly decreased in *Senp1* haploinsufficient mice, in comparison with WT mice, suggesting that the function of excitatory synapses was compromised in RSA neurons of *Senp1* haploinsufficient mice (Figure 4K-4M).

**Figure 4.**
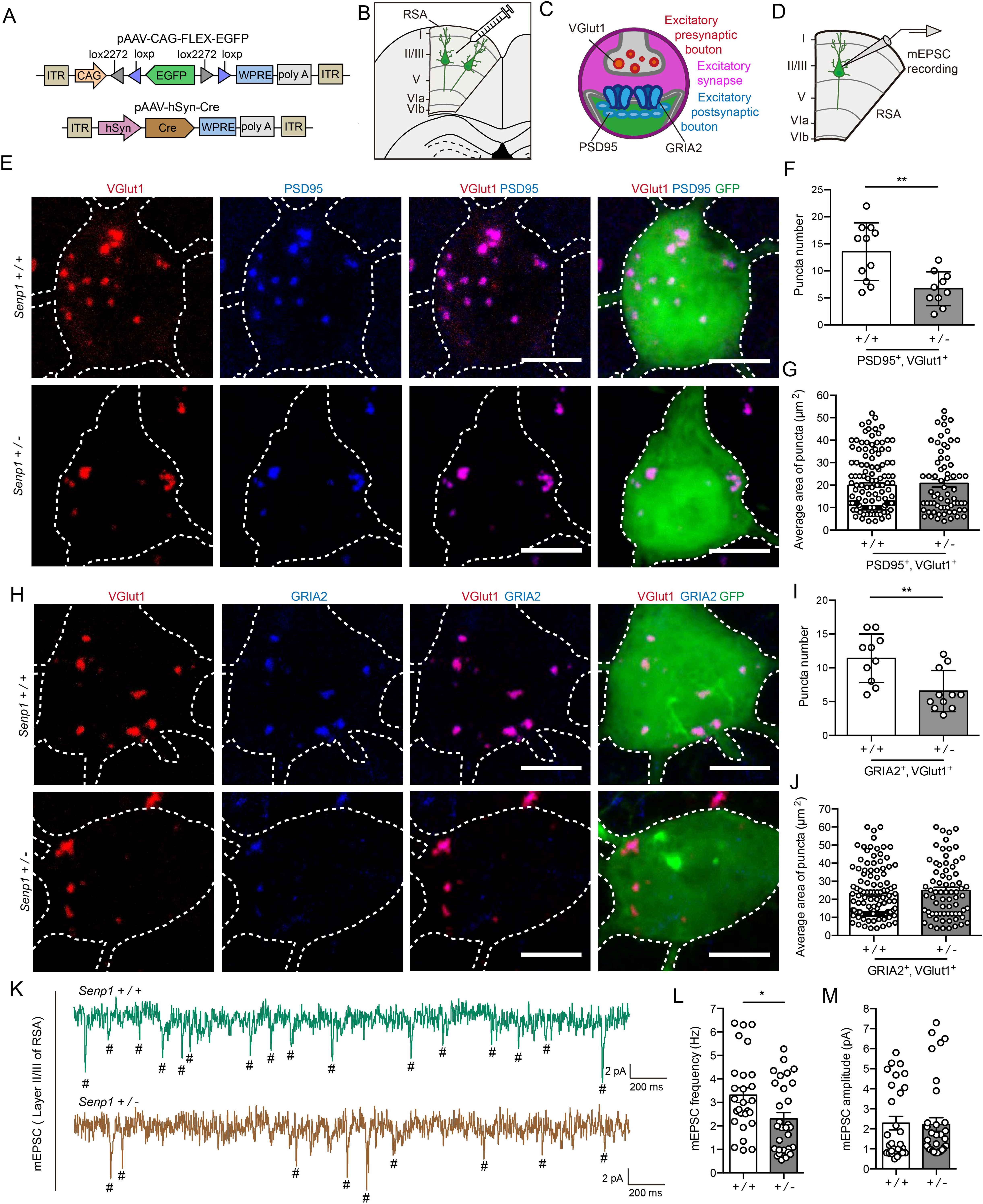
Decreased of excitatory synapses on pyramidal neuron correlates to restrained mEPSC frequency for *Senp1*-haploinsufficient mice. (A) Schematic illustration of AAV viral vector design strategy. (B) Schematic illustration for AAV injection to sparsely label pyramidal neurons in layer II/III of RSA. (C) Schematic illustration of immunofluorescence staining strategy to mark the excitatory synapse, stained for excitatory presynaptic marker, VGlut1 (red), and excitatory postsynaptic marker, PSD95 (blue), and excitatory postsynaptic marker, GRIA2 (blue). (D) Schematic illustration of brain slice recording for mEPSC of pyramidal neuron in RSA. (E) Representative immunofluorescence staining pictures for excitatory synapses on soma part of sparsely labeled pyramidal neuron in RSA, stained for VGlut1 (red), PSD95 (blue), and GFP (green). Scale bar, 60µm. (F) Quantification of puncta number of both PSD95 positive and VGlut1 positive bouton on the soma (neuron number n = 11, from 3 *Senp1 ^+/+^* mice; n = 10, from 3 *Senp1 ^+/-^* mice). (G) Quantification of average area for the puncta (≥ 4µm^2^) of both PSD95 positive and VGlut1 positive bouton on the soma (punctum number n = 139, from 11 neurons calculated in Figure 4F, for *Senp1 ^+/+^* mice; n = 67, from 10 neurons calculated in Figure 4F, for *Senp1 ^+/-^* mice). (H) Representative immunofluorescence staining pictures for excitatory synapses on soma part of sparsely labeled pyramidal neuron in RSA, stained for VGlut1 (red), GRIA2 (blue), and GFP (green). Scale bar, 60µm. (I) Quantification of puncta number of both GRIA2 and VGlut1 positive bouton on the soma (neuron number n = 10, from 3 *Senp1 ^+/+^* mice; n = 11, from 3 *Senp1 ^+/-^* mice). (J) Quantification of average area for the puncta (≥ 4µm^2^) of both GRIA2 positive and VGlut1 positive bouton on the soma (punctum n = 114, from 10 neurons calculated in Figure 4I, for *Senp1 ^+/+^* mice; n = 72, from 11 neurons calculated in Figure 4I, for *Senp1 ^+/-^* mice). (K) Representative mEPSC traces of pyramidal neurons in the layer II/III of RSA. (L) Quantification of mEPSC frequency for pyramidal neurons in the layer II/III of RSA (neuron number n = 27 neurons from 3 *Senp1 ^+/+^* mice; n = 30 from 3 *Senp1 ^+/-^* mice). (M) Quantification of mEPSC amplitude for pyramidal neurons in the layer II/III of RSA (neuron number n = 27 from 3 *Senp1 ^+/+^* mice; n = 30 from 3 *Senp1 ^+/-^* mice). * *p* < 0.05, ** *p* < 0.01. Bars represent means ± SD for Figure 4F and 4I; bars represent means ± SEM for Figure 4G, 4J, 4L and 4M.

It is worthy to note that the orchestrating alteration of inhibitory and excitatory synaptic transmission found in this study echoed a previous finding, in which there were also increased inhibitory synaptic transmission and decreased synaptic transmission in *Cntnap3* knockout mice, another autism mouse model that was previously reported by our lab (Tong et al., 2019). These shared phenotypes in different autism mouse models strongly suggest that the co-regulation of inhibitory and excitatory synapse may be one of the key features of autism pathophysiology in the brain.

### Impaired dendritic growth and immature spine of the RSA neurons of *Senp1* haploinsufficient mice

To investigate the role of SENP1 in the neuronal development, we constructed short hairpin RNA (shRNA) against the mouse *Senp1* gene (Figure S4A, S4B). By transfecting shRNA against *Senp1* and control shRNA in mouse cortical neurons, we found that the dendritic complexity was dramatically decreased when the endogenous SENP1 is knocked down by shRNA, which could be fully rescued by re-introducing human SENP1 open reading frame (SENP1 ORF), but not human SENP1 ORF carrying the stop-gain mutation found in the ASD patient (SENP1 stop) (Figure S4C, S4D). When SENP1 ORF was overexpressed in mouse cortical neurons, the overall dendritic complexity was increased, suggesting that SENP1 indeed plays a critical role in regulating dendritic growth (Figure. S4C, S4D). Interestingly, we also found that there are much more filopodia appeared in the SENP1 knocked down neurons comparing to GFP-expressing neurons which could be rescued by SENP1 ORF expression, suggesting that deletion of SENP1 may lead to impairment of spine maturation (Figure S4C, S4E).

We next examined the dendritic morphology of the layer II/III neurons in the RSA cortex of *Senp1* haploinsufficient mice by sparse labeling methods described above (Figures 3A). We found that the basal dendrites of layer II/III neurons exhibited less complexity in *Senp1* haploinsufficient mice comparing to WT mice, whereas apical dendrites appeared to be more complex in *Senp1* haploinsufficient mice suggesting that basal and apical dendrites were regulated differentially *in vivo* (Figures 5A, 5B). When we examined the spine morphology of layer II/III neurons in the RSA cortex, we surprisingly found that although the total spine numbers increased in the basal and apical dendrites of layer II/III neurons in *Senp1* haploinsufficient mice (Figure 5C, 5D), there were much less spines in the mature form and more immature spines in layer II/III neurons of *Senp1* haploinsufficient mice (Figure 5E-5H), indicating that spine development in the RSA cortex was hindered by the haploinsufficiency of *Senp1*.

**Figure 5.**
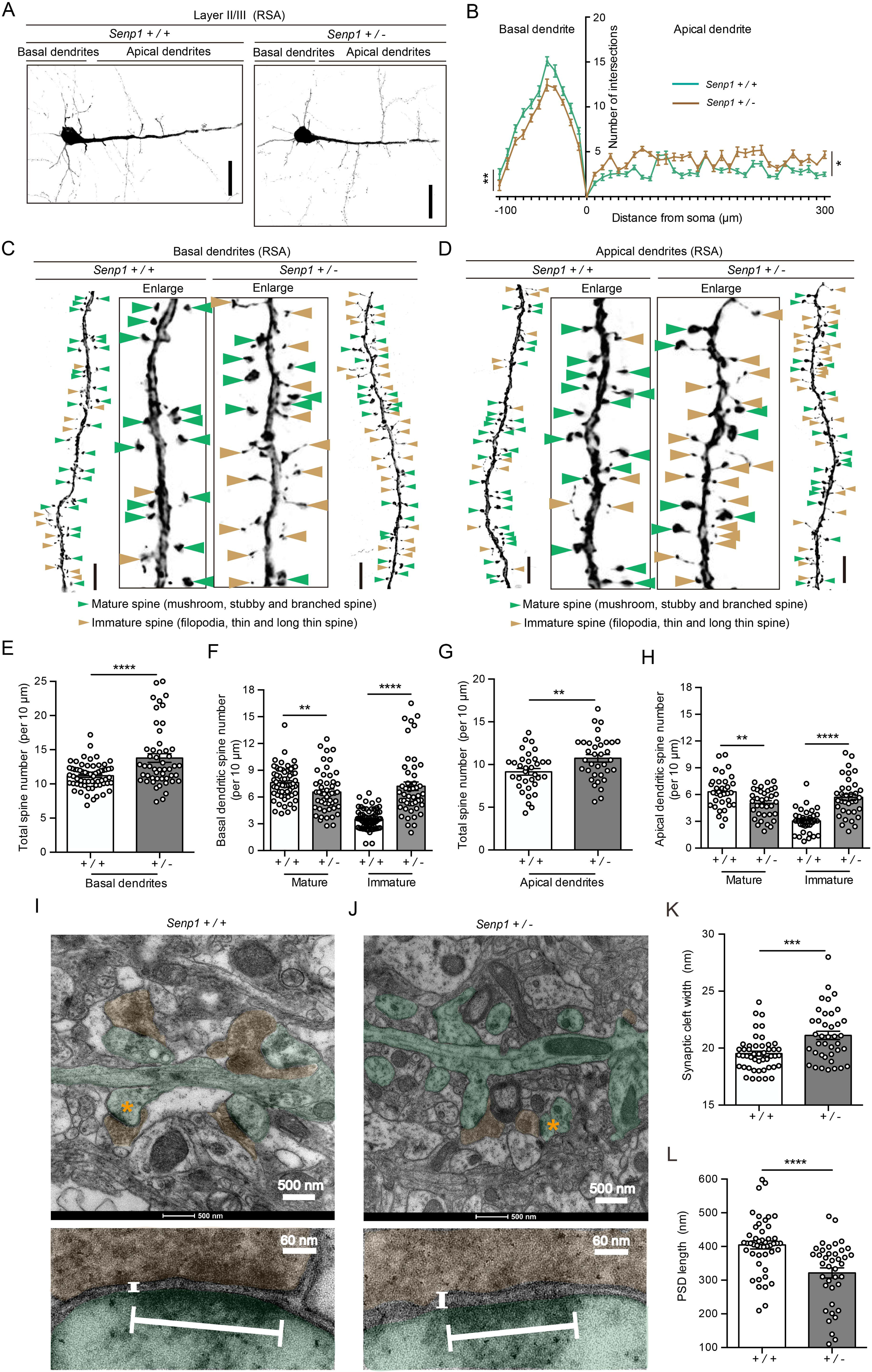
*Senp1*-haploinsufficient mice exhibit increased immature spine and abnormal ultrastructure of excitatory synapse. (A) Representative immunofluorescence staining pictures for sparsely labeled pyramidal neurons in layer II/III of RSA. Both basal dendrites and apical dendrites were illustrated. Scale bar, 100µm. (B) Quantification of Sholl-analysis calculated intersection number of branched dendrites according to distance from cell center (neuron number n = 12, from 3 mice for each genotype). (C) Representative immunofluorescence staining pictures for spines of basal dendritic part from sparsely labeled pyramidal neurons. Green arrowhead, mature spine (including mushroom, stubby and branched spine); brown arrowhead, immature spine (including filopodia, thin and long thin spine). Scale bar, 4µm. (D) Representative immunofluorescence staining pictures for spines of apical dendritic part from sparsely labeled pyramidal neurons. Scale bar, 4µm. (E) Quantification of spine density on basal dendrites. Spine numbers were counted on 10 μm long dendritic segments of secondary or tertiary dendrites. (62 segments collected from 14 neurons of 3 *Senp1 ^+/+^* mice; 50 segments collected from 10 neurons of 3 *Senp1 ^+/-^* mice). (F) Quantification of spine density for mature and immature spine on basal dendrites. Spine numbers were counted with the same way described as above (62 segments collected from 14 neurons of 3 *Senp1 ^+/+^* mice; 50 segments collected from 10 neurons of 3 *Senp1 ^+/-^* mice). (G) Quantification of spine density on apical dendrites. Spine numbers were counted with the same way described as above. (34 segments collected from 8 neurons of 3 *Senp1 ^+/+^* mice; 35 segments collected from 4 neurons of 3 *Senp1 ^+/-^* mice). (H) Quantification of spine density for mature and immature spine from apical dendrites. (34 segments collected from 8 neurons of 3 *Senp1 ^+/+^* mice; 35 segments collected from 4 neurons of 3 *Senp1 ^+/-^* mice). Representative transmission electron microscopy (TEM) pictures for microstructure of synapse in RSA of *Senp1 ^+/+^* mice (I) and *Senp1 ^+/-^* mice (J). (K) Quantification of synaptic cleft width (synapses number n=47, from 3 *Senp1 ^+/+^* mice; n=39 synapses, from 3 *Senp1 ^+/-^* mice). (L) Quantification of PSD length (synapses number n = 47, from 3 *Senp1 ^+/+^* mice; n=39 synapses, from 3 *Senp1 ^+/-^* mice). * *p* < 0.05, ** *p* <0.01, *** *p* < 0.001, **** *p* <0.0001. Bars represent means ± SEM. See also Figure S4.

We then examine the fine structure of excitatory synapse of the layer II/III neurons in RSA of *Senp1* haploinsufficient and WT mice with electron microscopy. Consistently, we found that the synaptic cleft appeared to be larger and the length of postsynaptic density (PSD) decreased in *Senp1* haploinsufficient mice (Figure 5I-5L), suggesting that SENP1 plays a critical role in promoting excitatory synapse maturation *in vivo*.

### Downregulation of FMRP in the RSA cortex of *Senp1* haploinsufficient mice

We would like to investigate the molecular mechanism by which SENP1 regulates synaptic development. Since SENP1 is one of the major SUMOylation deconjugating enzymes, we reasoned that SENP1 may regulate synapse development by conjugating small uniqitin-like modifier (SUMO) to candidate proteins, which performed critical functions in syanpse development. In a previous study, researchers identified that SENP1 proteins existed in the post-synaptic complex and implicated in brain disorders using mass spectrometry (Li et al., 2017). Among many synaptic proteins which are associated with SUMOylation modification, fragile X syndrome mental retardation protein (FMRP) was an intriguing candidate of SENP1-regulated deSUMOylation. The mutation of *FMR1*, the coding gene of FMRP, caused Fragile X syndrome (FXS), which shares autistic symptoms with ASD (Khayachi et al., 2018; Tang et al., 2018). It was reported that SUMOylation of FMRP was critical for its transporting mRNAs critical for synaptic functions in to synapses and release of mRNA from FMRP proteins(Khayachi et al., 2018). The SUMOylation and deSUMOylation is a dynamic process, dysregulation either of which will sure lead to compromise of proper functions of FMRP. Thus we set out to examine whether FMRP is dysregulated in the brain of *Senp1* haploinsufficient mice.

We first tested whether FMRP is expressed in the RSA of mouse brain. By immunostaining experiment using antibodies against SENP1 and FMRP, we found that FMRP closely colocalized with SENP1 in the RSA neurons of the mouse brain and was downregulated in *Senp1* haploinsufficient mice (Figure 6A, 6B). Moreover, we examined the expression of SENP1 and FMRP in the RSA cortex during mouse development and found that the expression levels of SENP1 and FMRP exhibit similar curve during mouse pre and post-natal brain development (Figure 6C-6E). Next we examined whether the expression of FMRP was regulated by SENP1 *in vitro* and *in vivo*. In the cultured mouse cortical neurons, we found that the level of FMRP protein significantly decreased after SENP1 knockdown (Figure 6F, 6G). We then performed Western blot on the tissue lysate collected from RSA region of *Senp1* haploinsufficient and WT mice (Figure 6B). We found that the protein level of FMRP in the RSA region was clearly downregulated in *Senp1* haploinsufficient mice comparing to its of WT mice (Figure 6H, 6I), strongly suggesting that FMRP was regulated by SENP1 in the RSA of the brain.

**Figure 6.**
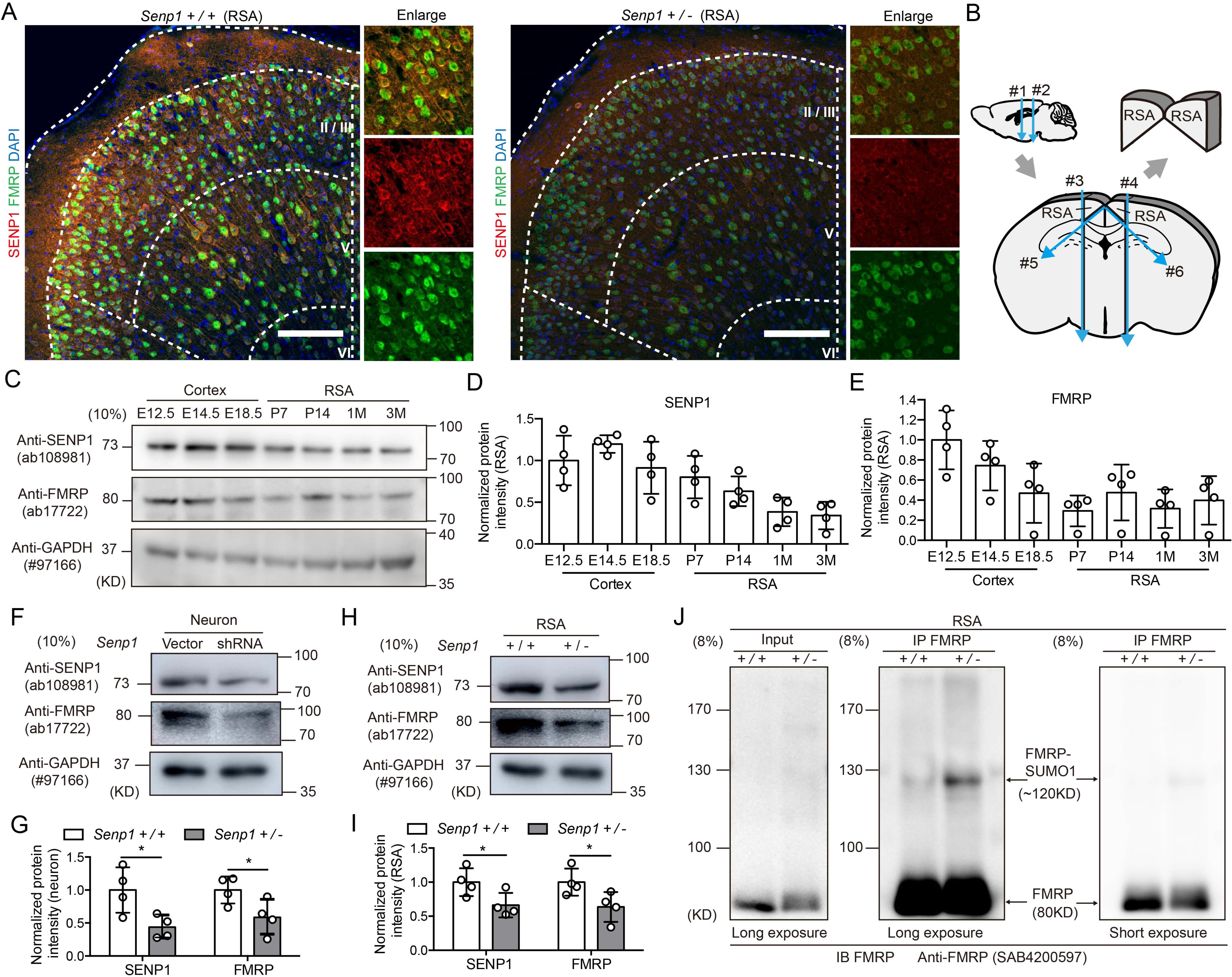
SENP1 regulates the expression of FMRP in RSA neurons via SUMOylation. (A) Representative coronal immunofluorescence images stained for SENP1 (red), FMRP (green) and DAPI (blue) in RSA of mice at 3 months of age. Scale bar, 200µm. (B) Schematic illustration for dissecting the RSA region from the mouse brain. (C) Representative western blot pictures illustrating protein levels of SENP1, FMRP and GAPDH in the whole cortex and the RSA region of *Senp1 ^+/+^* mice. E12.5-E18.5, embryonic day12.5-18.5; P7, P14, postnatal day 7 and14; 1M, 3M, 1 month and 3 months ago. Quantification of normalized protein levels of SENP1(D) and FMRP (E) for each time point (n = 4). (F) Representative western blot pictures showing protein levels of SENP1, FMRP and GAPDH for cultured cortical neurons. Neurons were electroporated with empty vector (vector), or mouse SENP1 shRNA (shRNA). (G) Quantification of normalized protein levels of SENP1 and FMRP (Data collected from 4 independent experiments). (H) Representative western blot pictures showing protein levels of SENP1, FMRP and GAPDH in the RSA region from *Senp1* ^+/+^ and *Senp1*^+/-^ mice. (I) Quantification of normalized SENP1 and FMRP protein intensity (n = 4, for each genotype). (J) Representative immunoblot with anti-FMRP antibody using RSA extracts of 3 months old mouse immunoprecipitated with anti-FMRP antibody or control IgG. * *p* < 0.05. Bars represent means ± SD.

If SENP1 indeed plays a critical role in removing the SUMO groups from FMRP protein, the SUMOylation level of FMRP in the brain of *Senp1* haploinsufficient mice may elevate. To determine the level of FMRP SUMOylation in the RSA region, we collected the RSA tissue lysate from *Senp1* haploinsufficient and WT mice, then performed immunoprecipitation assay using antibody against FMRP (Figure 6B). We found that in the RSA lysate from *Senp1* haploinsufficient mice, there is clearly an up-shifting band with FMRP positive signals, but not in WT mice (Figure 6J), suggesting that FMRP-SUMO signals are enriched in the RSA region of *Senp1* haploinsufficient mice. Together, these results suggest that SENP1 regulates the SUMOylation of FMRP in the brain, hereby over SUMOylation of FMRP protein in the brain of *Senp1* haploinsufficient mice may lead to its degradation (Khayachi et al., 2018).

### Restoring the expression of SENP1 or FMRP in RSA rescue autistic-like behaviors of *Senp1* haploinsufficient mice

Finally, we wonder whether restoration of SENP1 or FMRP in RSA could rescue the autistic-like behaviors of *Senp1* haploinsufficient mice. We injected AAVs harboring control vector, SENP1 or FMRP cDNA bi-laterally in RSA of *Senp1* haploinsufficient mice at 2 months of age and examined behaviors one month later (Figure 7A).

**Figure 7.**
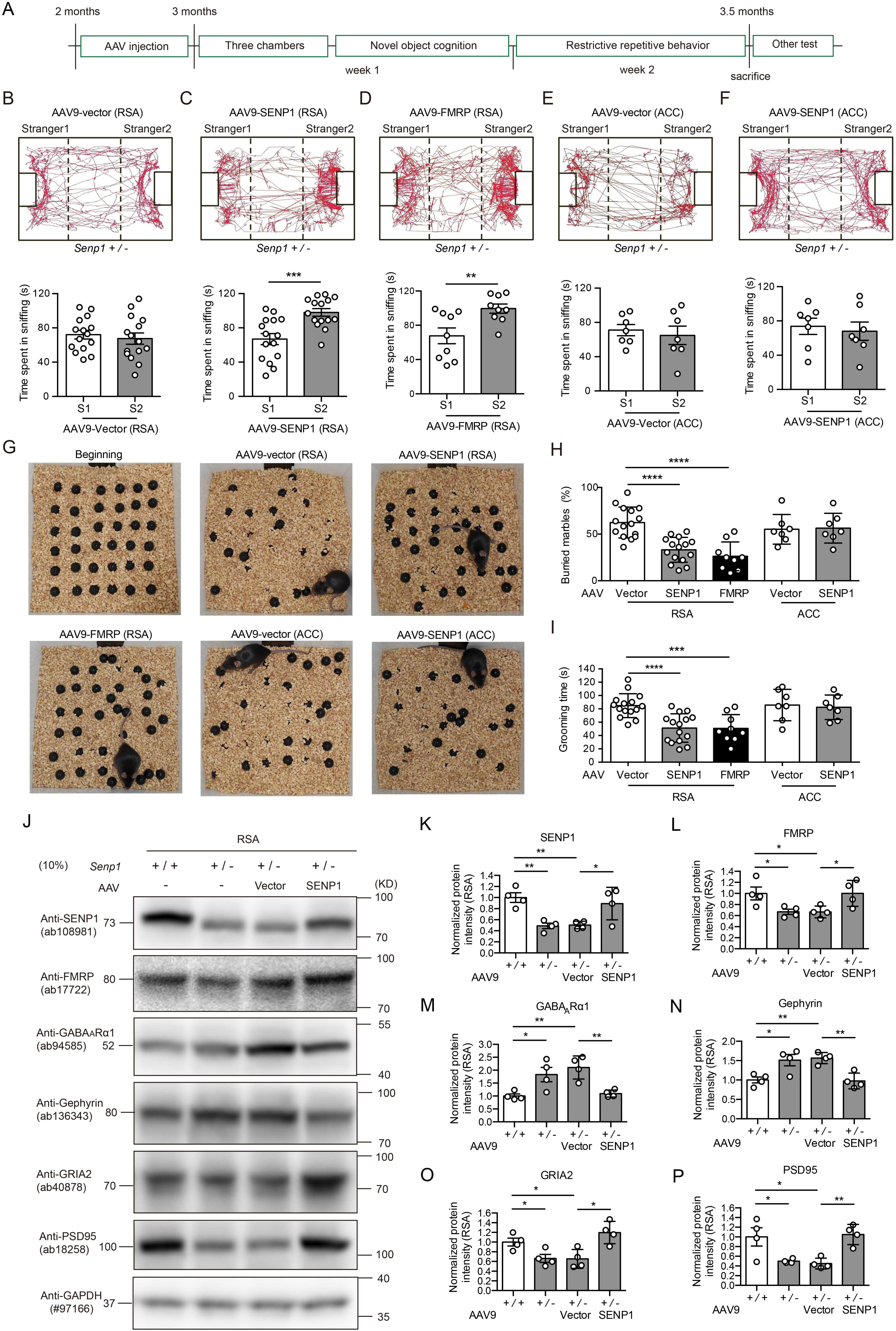
AAV delivery of SENP1 and FMRP in RSA region rescues autistic behaviors of *Senp1*-haploinsufficient mice. (A) Strategy of behavioral tests after AAV injection for *Senp1 ^+/-^* mice. (B) Representative locomotor traces of mice in the three-chamber test for social novelty (upper panel) and quantification of time mice spent in sniffing familiar and novel partners (lower panel). *Senp1 ^+/-^* mice were injected with AAV9-vector in RSA (n = 15). (C) Representative locomotor traces of mice in the three-chamber test for social novelty (upper panel) and quantification of time mice spent in sniffing familiar and novel partners (lower panel). *Senp1 ^+/-^* mice were injected with AAV9-SENP1 in RSA (n = 15). (D) Representative locomotor traces of mice in the three-chamber test for social novelty (upper panel) and quantification of time mice spent in sniffing familiar and novel partners (lower panel). *Senp1 ^+/-^* mice were injected with AAV9-FMRP in RSA (n = 9). (E) Representative locomotor traces of mice in the three-chamber test for social novelty (upper panel) and quantification of time mice spent in sniffing familiar and novel partners (lower panel). *Senp1 ^+/-^* mice were injected with AAV9-vector in ACC (n = 7). (F) Representative locomotor traces of mice in three-chamber test for social novelty (upper panel) and quantification of time mice spent in sniffing familiar and novel partners. (lower panel) *Senp1 ^+/-^* mice were injected with AAV9-SENP1 in ACC (n = 7). (G) Representative figures showing from beginning to the end of the marble burying test for *Senp1 +/-* mice after AAV injection. (H) Quantification of percentage of buried marbles within 10 min. *Senp1 ^+/-^* mice injected with various AAVs (AAV9-vector in RSA n = 15; AAV9-SENP1 in RSA n = 15; AAV9-FMRP in RSA n=9; AAV9-vector in ACC n = 7; AAV9-SENP1 in ACC n = 7). (I) Quantification of grooming time within 30 min. *Senp1 ^+/-^* mice injected with various AAVs (AAV9-vector in RSA n = 15; AAV9-SENP1 in RSA n = 15; AAV9-FMRP in RSA n = 9; AAV9-vector in ACC n = 7; AAV9-SENP1 in ACC n = 7). (J) Representative western blot pictures illustrating protein intensity of SENP1, FMRP, GABA_A_Rα1, Gephyrin, GRIA2, PSD95 and GAPDH for RSA tissues of mice with or without injection of AAV-vector or AAV-SENP1. Quantification of normalized protein intensity of SENP1 (K), FMRP (L), GABA_A_Rα1 (M), Gephyrin (N), GRIA2 (O) and PSD95 (P) (n = 4 for each genotype, with or without injection of AAV9-vector or AAV9-SENP1 in RSA). * *p* < 0.05, ** *p* <0.01, *** *p* < 0.001, **** *p* <0.0001. Bars represent means ± SD.

We found that injection of control AAV (AAV-vector) did not change the deficits in social novelty test of *Senp1* haploinsufficient mice (Figure 7B), whereas injection of AAV-SENP1 into the RSA of *Senp1* haploinsufficient mice significantly increased the time *Senp1* haploinsufficient mice spent in sniffing novel partners (Figure 7C), suggesting that the SENP1 in the RSA is crucial for social behaviors in mouse. Similarly, injection of AAV-FMRP in the RSA region of *Senp1* haploinsufficient mice also alleviated the social deficits (Figure 7D).

To further examine the specificity of RSA in social behaviors, we asked whether injection of AAV-SENP1 into the anterior cingulate cortex (ACC), a well-known region directly connected to RSA and also implicated in autistic behaviors (Guo et al., 2019). Surprisingly, we found that injection of AAV-SENP1 into the ACC had no effects on social behaviors of *Senp1* haploinsufficient mice (Figure 7E, 7F). Consistently, we found that the stereotypic and repetitive behaviors of *Senp1* haploinsufficient mice were fully rescued by injection of AAV-SENP or AAV-FMRP into RSA, but not ACC (Figure 7G-7I).

Next, we wondered whether synaptic abnormalities in the RSA cortex of *Senp1* haploinsufficient mice may be rescued by genetic manipulations. First, we examined levels of various synaptic protein with biochemical methods. We collected tissue lysates from RSA regions of WT and *Senp1* haploinsufficient mice with or without AAV injection (Figure 7J). Using Western blots with antibodies against various proteins, we found that re-introducing SENP1 in the RSA of *Senp1* haploinsufficient mice restored the increased level of the GABA_A_R α1 subunit and gephyrin to the normal level as WT mice (Figure 7K-7N), as well as the decreased level of GRIA2 and PSD95 back to levels in the WT mice (Figure 7O, 7P). Consistently, injection of AAV-FMRP was also able to rescue the biochemical defects in the *Senp1* haploinsufficient mice (Figure S5A-S5G).

### Synaptic defects in the RSA of *Senp1* haploinsufficient mice were rescued by restoration of SENP1 or FMRP

We then examined whether the structure and function of RSA neurons in *Senp1* haploinsufficient mice could be rescued by re-introducing SENP1 or FMRP. By the sparse labeling methods used above, we first measured the inhibitory and excitatory synapses in RSA layer II/III neurons of *Senp1* haploinsufficient mice injected with AAVs expressing control vector, SENP1 or FMRP, perspectively (Figure 8A, 8B). We found that re-introducing SENP1 or FMRP into the RSA of *Senp1* haploinsufficient mice significantly decreased the abnormal high inhibitory synapse number, represented by punctum co-labelled with Gephyrin and VGAT (Figure 8C). By performing whole-cell patch clamp experiments, we found that mIPSC frequency also accordingly decreased in the layer II/III RSA neurons in *Senp1* haploinsufficient mice injected with AAV-SENP1 or AAV-FMRP, comparing with AAV-vector injecting mice (Figure 8D, 8E).

**Figure 8.**
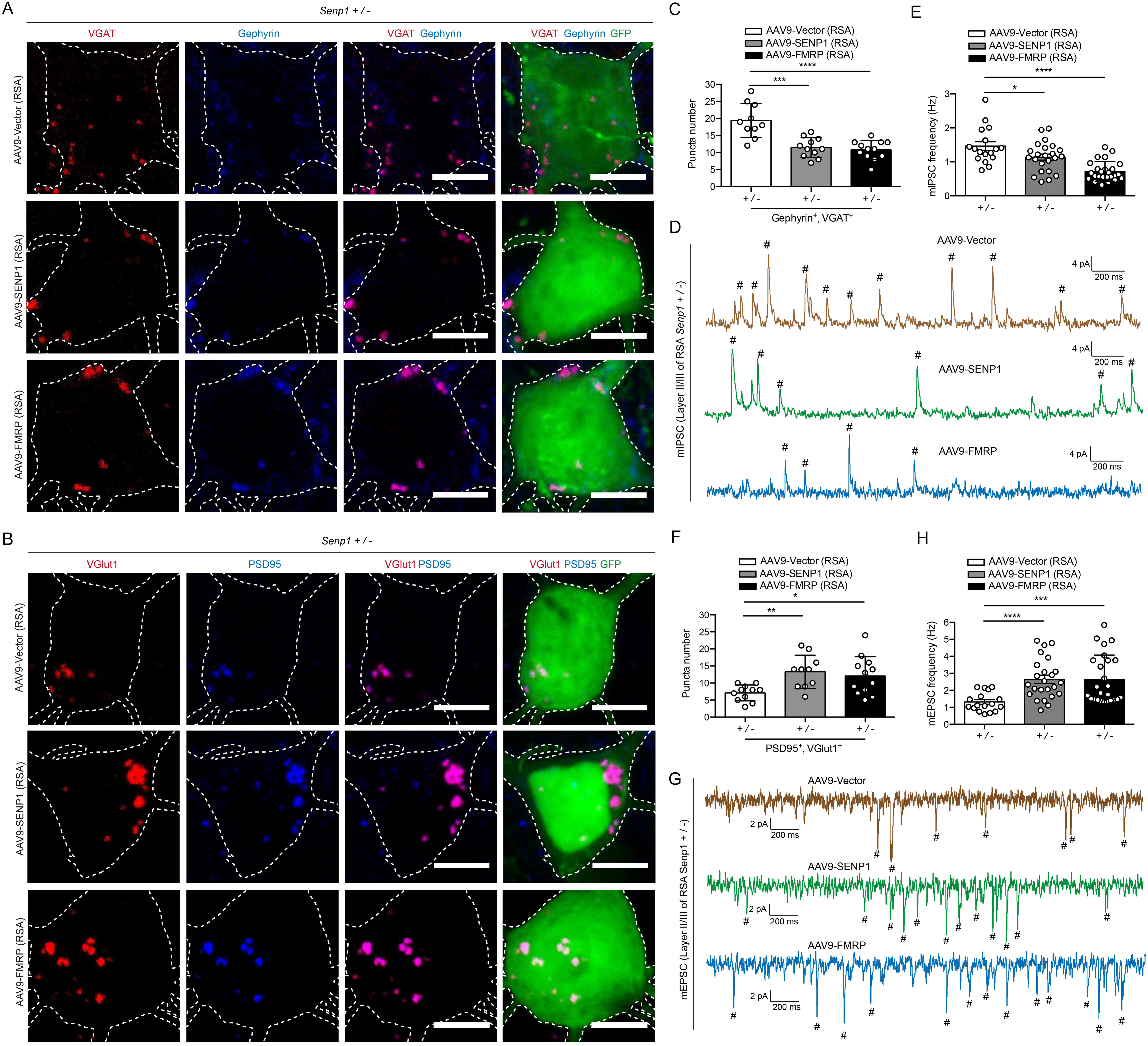
AAV delivery of SENP1 and FMRP in RSA region rescues abnormal inhibitory and excitatory synapse in pyramidal neuron. (A) Representative immunofluorescence images for inhibitory synapses on sparsely labeled pyramidal neurons in RSA, VGAT (red), Gephyrin (blue), and GFP (green). Scale bar, 60µm. (B) Representative immunofluorescence images for excitatory synapses on sparsely labeled pyramidal neurons in RSA, stained for VGlut1 (red), PSD95 (blue), and GFP (green). Scale bar, 60µm. (C) Quantification of puncta number of both Gephyrin and VGAT positive bouton on the soma of layer II/III neurons in RSA (AAV9-vector, n=10 neurons; AAV9-SENP1, n=11 neurons; AAV9-FMRP, n=12 neurons). (D) Representative mIPSC traces of pyramidal neurons in the layer II/III of RSA. (E) Quantification of mIPSC frequency for pyramidal neurons in the layer II/III of RSA (AAV9-vector, n=17 neurons; AAV9-SENP1, n=23 neurons; AAV9-FMRP, n=25 neurons). (F) Quantification of puncta number of both PSD95 positive and VGlut1 positive bouton on the soma of RSA layer II/III neurons (AAV9-vector, n=11 neurons; AAV9-SENP1, n=10 neurons; AAV9-FMRP, n=12 neurons). (G) Representative mEPSC traces of pyramidal neurons in the layer II/III of RSA. (H) Quantification of mIPSC frequency for pyramidal neurons in the layer II/III of RSA (AAV9-vector, n=17 neurons; AAV9-SENP1, n=24 neurons; AAV9-FMRP, n=25 neurons). * *p* < 0.05, ** *p* <0.01, *** *p* < 0.001, **** *p* <0.0001. Bars represent means ± SD for Figure 8C and Figure 8F; bars represent means ± SEM for Figure 8E and Figure 8H. Data collected from 3 *Senp1 ^+/-^* mice injected with AAVs.

Interestingly, re-introducing SENP1 or FMRP increased the number of excitatory synapse in the layer II/III RSA neurons of *Senp1* haploinsufficient mice, measured by punctum with PSD95 and VGlut1 signals (Figure 8B, 8F). Consistently, mEPSCs frequency of RSA layer II/III neuron in *Senp1* haploinsufficient mice injected with AAV-SENP1 or AAV-FMRP significantly increased, comparing to mice injected with AAV-vector (Figure 8G, 8H). Together, these data indicate that the SENP1-FMRP axis regulates the balance of inhibitory and excitatory synapse in the RSA region of the mouse brain.

## Discussion

Genomic studies using whole-exome sequencing technology for large ASD cohorts since 2011 have proven to be very fruitful for identification of ASD-causing genetic mutations (Iossifov et al., 2012; Neale et al., 2012; O’Roak et al., 2011; O’Roak et al., 2012; Sanders et al., 2012). Besides the classic dominant and recessive mutations, the *de novo* mutations taking place usually at male germ cells are found to play a critical role in contributing to the etiology of ASD. The gene-disrupting mutations (usually referred as likely gene disrupting—LGD) are more often to be discovered in the ASD probands, comparing to their unaffected siblings (Iossifov et al., 2014). Although the overall contribution of *de novo* LGD mutations may account for around 10% of genetic causes of ASD, they open a venue for us to understand the mechanism by which the brain receives and processes social signals and perform social interaction behaviors accordingly. 50-70% of ASD cases would have comorbidity of intellectual disability or developmental delay (Lord et al., 2020). Searching for genetic causes in ASD patients without comorbidity of developmental delay may lead to discovery of genes specific for social behaviors in mammals (Satterstrom et al., 2020). There certainly is a possibility that there are not any genes specific for social behaviors, since different penetrance of genetic mutations on individual may lead to various extent of symptoms (Myers et al., 2020). However, it is still the consensus that focusing on the genetic mutations primarily affecting core symptoms of autism may shed new light on the understanding of neural circuits underlying social interaction behaviors (Myers et al., 2020).

In this work, we found that in the *Senp1* haploinsufficient mice, mimicking the human ASD patient, both inhibitory and excitatory synaptic transmission in the RSA region of retrosplenial cortex was specifically affected, which causally lead to deficits in social novelty tests and could be rescued by re-introducing SENP1 or its downstream molecule FMRP during adulthood (Figure 9A, 9B). This data suggests that the RSA region plays a critical role in regulating social behaviors in the mouse brain. From previous works, the retrosplenial cortex is implicated in the top-down control of sensorimotor information integration, as well as playing a role in coordinated oscillation with hippocampal neurons (Mao et al., 2018). Together with the recent finding that ketamine treatment in mice specifically activated the RSC neurons in mice and lead to dissociative-like symptoms, it is intriguing to hypothesize that the RSC region may play a critical role in orchestrating sensory information from primary cortices to higher centers, in which abnormal neural activity caused either by acute ketamine treatment or developmental defects due to lack of SENP1, would lead to various behavioral deficits, including social interactions (Vesuna et al., 2020).

**Figure 9.**
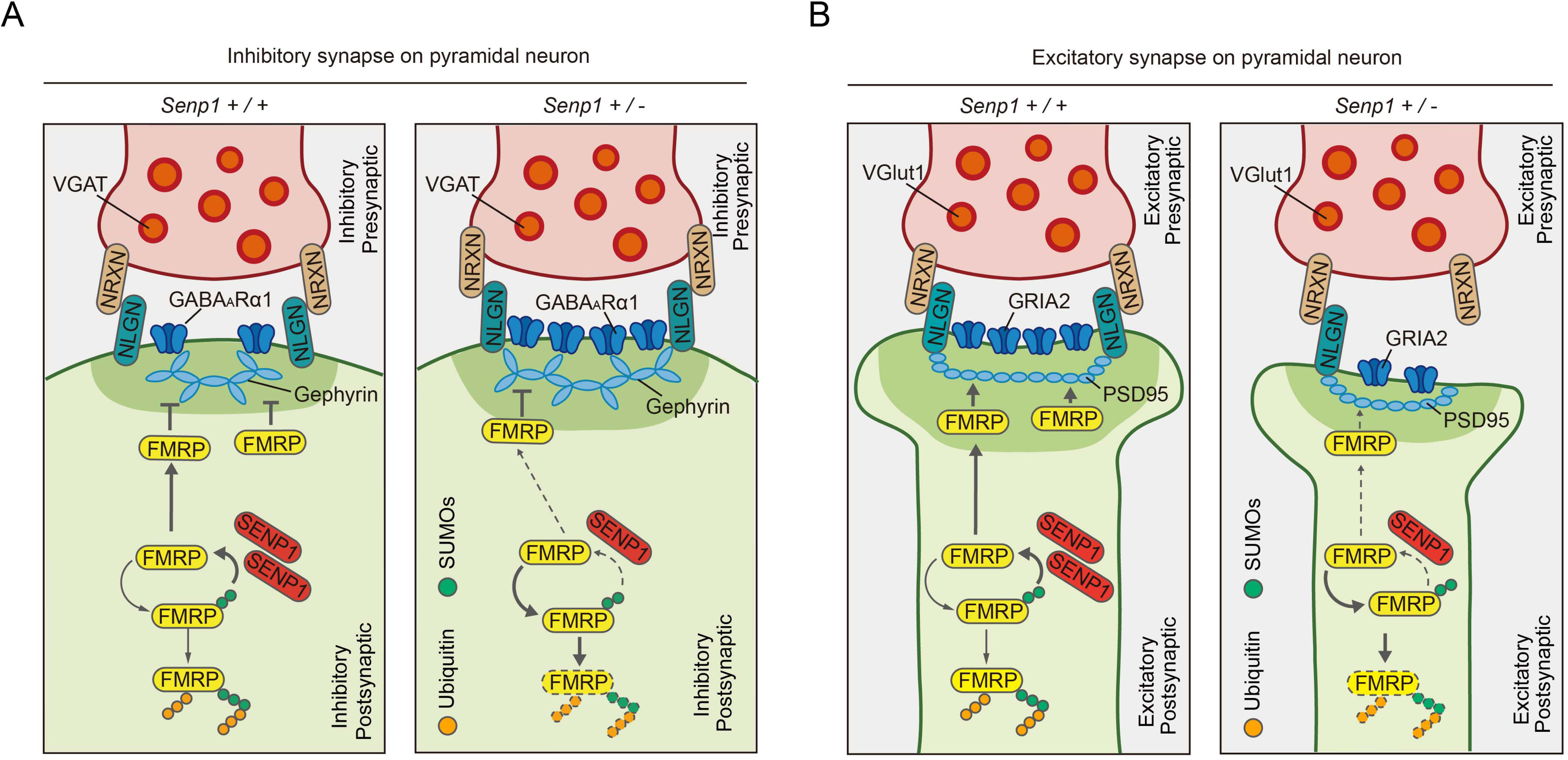
Schematic illustration for alterations of both inhibitory and excitatory synapses in *Senp1* haploinsufficient mice. (A) Schematic illustration for inhibitory synapses in *Senp1 ^+/+^* and *Senp1 ^+/-^* mice. NLGN, Neuroligin; NRXN, Neurexin. (B) Schematic illustration for excitatory synapses in *Senp1 ^+/+^* and *Senp1 ^+/-^* mice. NLGN, Neuroligin; NRXN, Neurexin.

Although ASD is a developmental disorder, in which genetic defects usually affect the brain development from early stages, it is widely recognized that autistic-like phenotypes in autism mouse models, such as *Mecp2* mutant or duplication mice, could be rescued with genetic methods (Guy et al., 2007; Sztainberg et al., 2015) or by gene editing tools (Qiu, 2018; Sun et al., 2020a; Yu et al., 2020) during adulthood. These works indicate that the neural circuits governing social behaviors still remain plastic after brain is fully grown, thus identification of which will provide the possibility of manipulation impaired social behaviors in adolescent or adult ASD patients.

The neural mechanisms by which RSA regulates social behaviors is intriguing. The role of retrosplenial cortex is implicated in integration of sensory information from primary cortices. The sensorimotor problems are prevalent in ASD patients which also become one of major targets for recovery therapies (sensory integrative training). One of primary hypothesis for etiology of ASD is that abnormal development may disrupt the sensory integration and in turn compromised social interaction behaviors. Although the brain processes many kinds of sensory inputs during everyday life, whether there is a specific neural circuit responsible for process and relay social-related sensory inputs is unknown. Thus, our current work leads to the hypothesis framework that RSA may be able to extract the social information received from primary sensory cortices and present them to higher centers. Further works mapping the neural circuits upstream and downstream of RSA neurons would illustrate the comprehensive picture of social information processing in the brain. Furthermore, the availability of non-human primate models for autism may facilitate the translational efforts in which non-invasive or invasive neural modulation methods targeting RSA would be applied and yield valuable insights for intervention of ASD, prior to clinical practices in patients (Liu et al., 2016; Zhou et al., 2019)

---

**Supplemental Figure 1-5, supplemental table 1, methods and materials are with online version of this paper.**

## Acknowledgements

We thank Drs. Xiaohong Xu, Ji Hu, Xiaoke Chen, Yu Fu and the Neuroscience Pioneering Club for valuable comments and Qian Hu for excellent technical assistance. This work was supported by grants from the NSFC Grants (#31625013, #81941015, #82001211, #82021001, #81761128035, #81930095), Strategic Priority Research Program of the Chinese Academy of Sciences (XDBS01060200), Program of Shanghai Academic Research Leader, the Open Large Infrastructure Research of Chinese Academy of Sciences, and the Shanghai Municipal Science and Technology Major Project (#2018SHZDZX05), CPSF-CAS Joint Foundation for Excellent Postdoctoral Fellows (2017LH036) and China Postdoctoral Science Foundation (2017M620173). Shanghai Municipal Commission of Health and Family Planning (#GWV-10.1-XK07, #2020CXJQ01), Shanghai Committee of Science and Technology (#2018SHZDZX01, #19410713500, #16JC1420501), Guangdong Key Scientific and Technological Project (#2018B030335001)

## Author contributions

K.Y. and Z.Q. designed the experiments, analyzed the data, and wrote the manuscript. K.Y. performed biochemistry, immunohistochemistry, and animal behavioral analysis experiments. Y.-H.S. performed stereotactic injections for immunohistochemistry experiments and electrophysiological recordings with the assistance of L.F. and H.-T.X.. X.-J.D. and J.-H.Y performed autism diagnosis under the supervision of F.L. J.-C.W performed whole-exome sequencing data analysis. Y.-F.Z. and Y.-T.Y. performed behavioral testing. S.-F.S. performed cell culture and tissue dissection. R.-Q.W., C.-H.Z., Y.-T.L., Z.-L.C. and Y.-Z.W performed Western blot and immunofluorescence. J.-K.C. provided *Senp1-*haploinsufficinet mice.

