## supplemental figures for "SENP1 in the retrosplenial agranular cortex regulates core autistic-like symptoms in mice"

### Table S1

### Figure S1-S5

### Table S1 Clinical diagnosis of ASD patient carrying *SENP1* mutation

**Supplemental Table 1. Yang et al.**

| **Age** | | 26.5 months | Note |
|---|---|---|---|
| **ADOS (scores)** | Communication | 8 | ASD (>2) |
| | Social interaction | 14 | ASD (>4) |
| **CARS (scores)** | | 39 | Severe (31-45) |
| **Gesell development scale (Developmental Quotient score)** | DQ at motor domain | 87 | Equal to 23 months (developmental age) |
| | DQ at fine motor domain | 113 | Equal to 30 months (developmental age) |
| | DQ at adoptive behaviors domain | 100 | Equal to 26.6 months (developmental age) |
| | DQ at language development domain | 68 | Equal to 18 months (developmental age) |
| | DQ at personal social development domain | 63 | Equal to 16.8 months (developmental age) |
| | DQ | 86.2 | Equal to 22.9 months (developmental age) |

### Supplemental Figures

Kan Yang et al

Figure S1

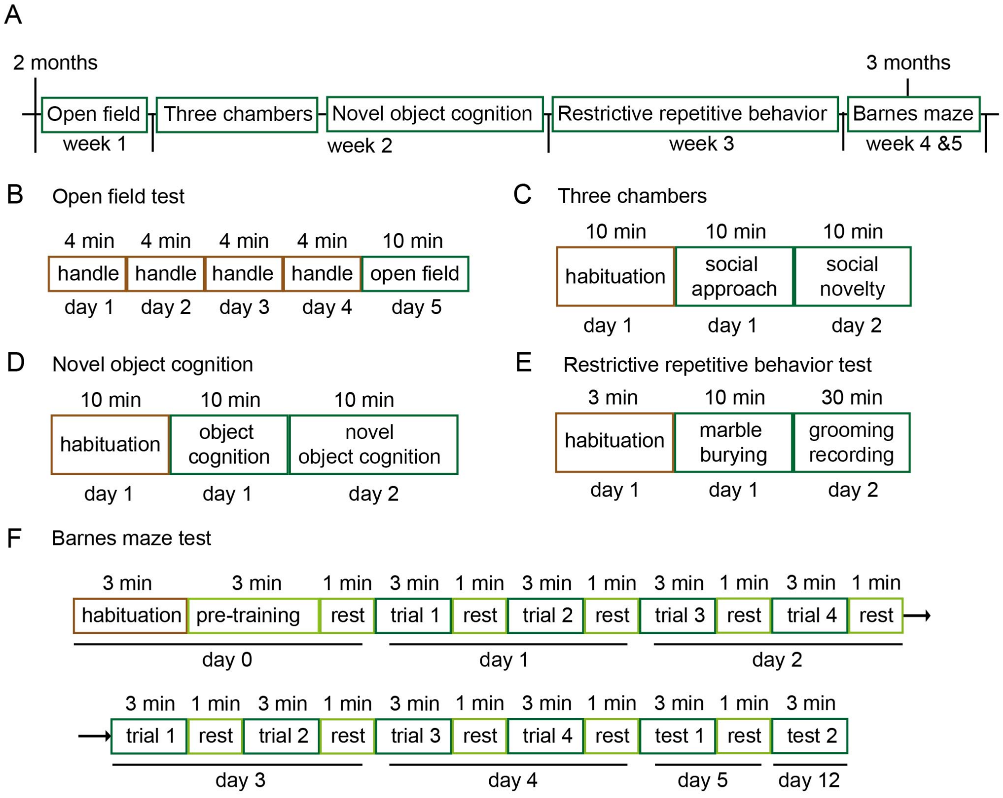


**Figure S1 Workflow for all behavioral test.**

(A) Strategy of behavioral tests. *Senp1*-haploinsufficient mice and WT littermates were applied to tests at 2 months of age. Only male mice were used in this study.

(B) Workflow for the open field test. Before the test, mice were gently handled for 4 consequent days.

(C) Workflow for three-chamber social behavioral tests, including social approach and social novelty.

(D) Workflow for three-chamber object cognition and novel object cognition tests.

(E) Workflow for restrictive repetitive behavior tests, including marble burying and grooming recording.

(F) Workflow for Barnes maze test. Pre-training (day 0) was applied to ensure the mice entry the target hole, before training stage (from day1 to day 5) and formal test (day 12).

Except for the open field test, mice were introduced with 10 min habituation in novel environment for all tests. See also Figure 1 and Figure S2.

Kan Yang et al

Figure S2

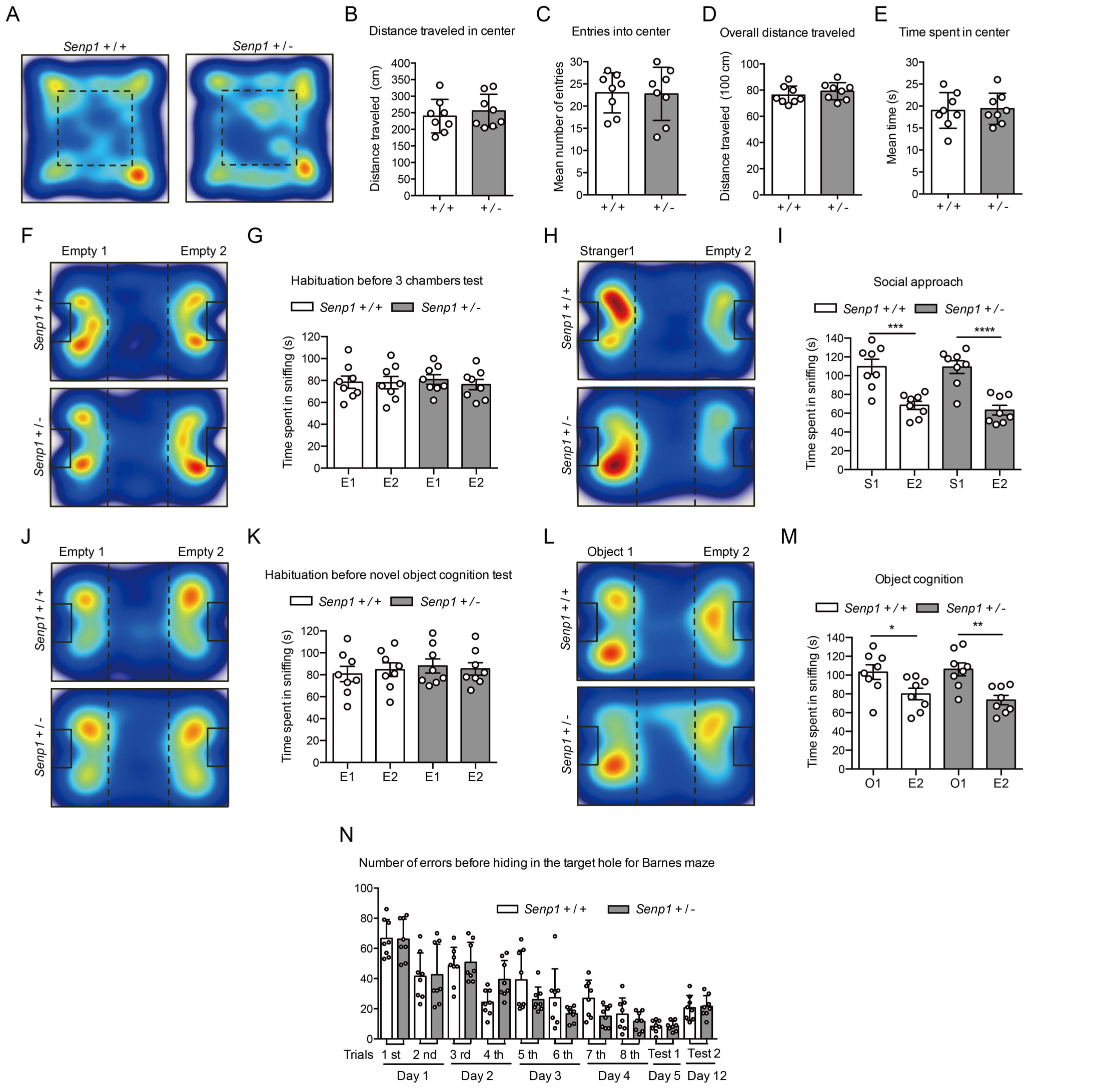


**Figure S2 *Senp1*-haploinsufficient mice exhibit no anxiety phenotypes, normal social approach, normal object cognition behaviors and normal spatial learning behaviors.**

(A) Representative heat map illustrating motion track of mice in the open field test.

(B) Quantification of distance mice traveled in the center zoom within 10 min (n = 8, for each genotype).

(C) Quantification of mean numbers mice entries into the center zoom within 10 min (n = 8, for each genotype).

(D) Quantification of overall distance mice traveled within 10 min (n = 8, for each genotype).

(E) Quantification of mean time mice spent in the center zoom within 10 min (n = 8, for each genotype).

(F) Representative heat map illustrating motion track of mice in habituation before three-chamber social approach test.

(G) Quantification of time mice spent in sniffing two empty cages on each side (n = 8, for each genotype).

(H) Representative heat map illustrating motion track of mice in three-chamber social approach test.

(I) Quantification of time mice spent in sniffing empty cages and partner (n = 8, for each genotype).

(J) Representative heat map illustrating motion track of mice in habituation before three-chamber object cognition test.

(K) Quantification of time mice spent in sniffing two empty cages on each side (n = 8, for each genotype).

(L) Representative heat map illustrating motion track of mice in three-chamber object cognition test.

(M) Quantification of time mice spent in sniffing empty cages and object (n = 8, for each genotype).

(N) Quantification of number of error before hiding in the target hole for Barnes maze (n = 8, for each genotype).

* $p < 0.05$, ** $p < 0.01$, *** $p < 0.001$, **** $p < 0.0001$. Bars represent means ± SD. See also Figure 1 and Figure S1.

Kan Yang et al

Figure S3

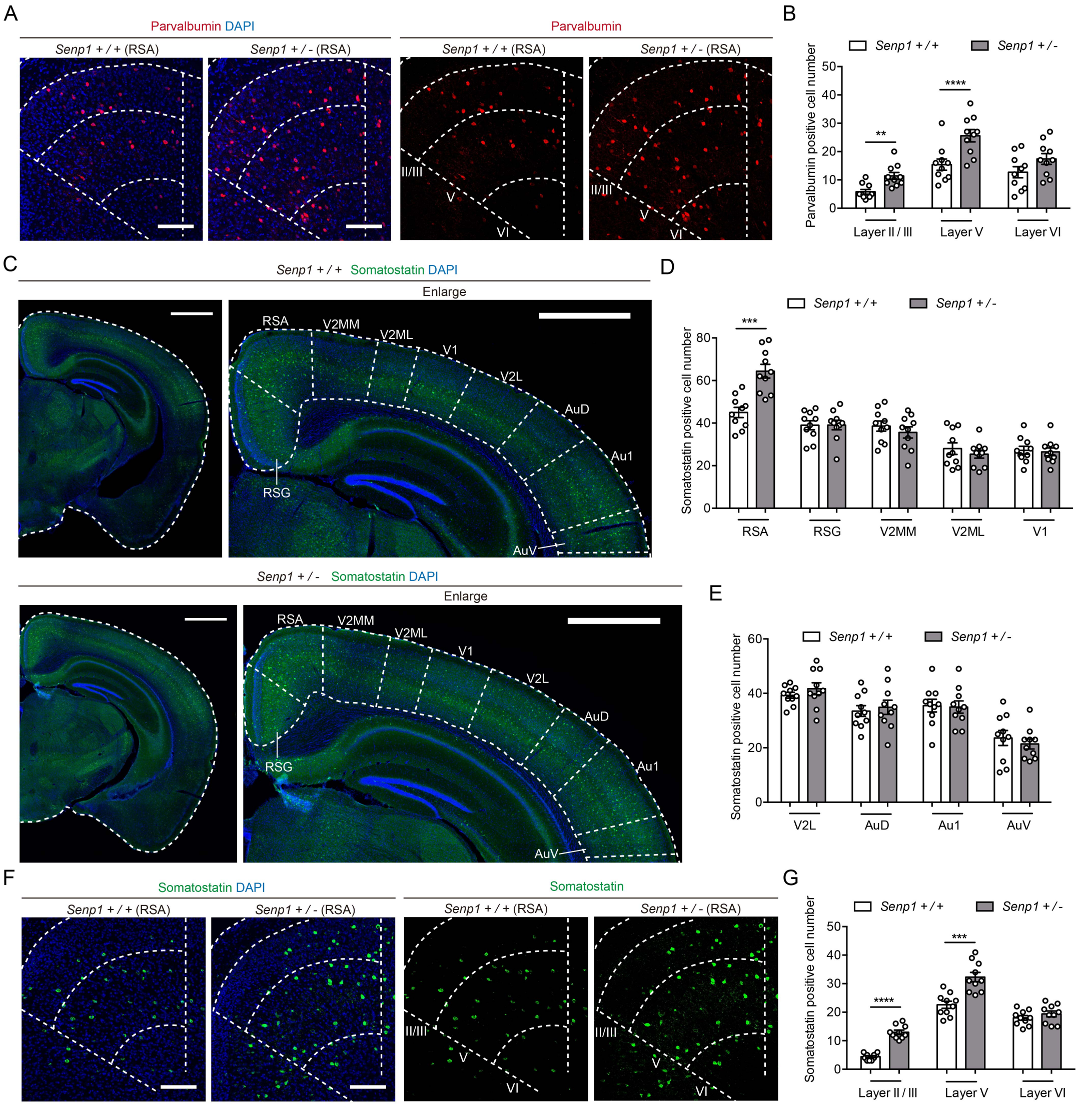


**Figure S3 *Senp1*-haploinsufficient mice exhibit RSA-specific increase of somatostatin positive interneuron and increase of parvalbumin and somatostatin positive interneuron within layer II/III and layer V.**

(A) Representative coronal immunofluorescence staining sections of retrosplenial agranular cortex stained for parvalbumin (red) and DAPI (blue) at 3 months. II/III, layer II/III; V, layer V; VI, layer VI. Scale bar, 200μm.

(B) Quantification of parvalbumin positive cell number within each layer (n = 3, for each genotype).

(C) Representative coronal immunofluorescence staining sections of brain stained for Somatostatin (green) and DAPI (blue) at 3 months. RSA, retrosplenial agranular cortex; RSG, retrosplenial granular cortex; V2MM, secondary visual cortex, mediomedial area; V2ML, secondary visual cortex, mediolateral area; V1, primary visual cortex; V2L, secondary visual cortex, lateral area; AuD, secondary auditory cortex, dorsal area; Au1, primary auditory cortex; AuV, secondary auditory cortex, ventral area. Scale bar, 1mm.

(D) Quantification of somatostatin positive cell number within each region (n = 3, for each genotype).

(E) Quantification of somatostatin positive cell number within each region (n = 3, for each genotype).

(F) Representative coronal immunofluorescence staining sections of retrosplenial agranular cortex stained for somatostatin (red) and DAPI (blue) at 3 months. II/III, layer II/III; V, layer V; VI, layer VI. Scale bar, 200μm.

(G) Quantification of somatostatin positive cell number within each layer (n = 3, for each genotype).

** $p < 0.01$, *** $p < 0.001$, **** $p < 0.0001$. n is the number of mice used in each experiment. Bars represent means ± SD.

Kan Yang et al

Figure S4

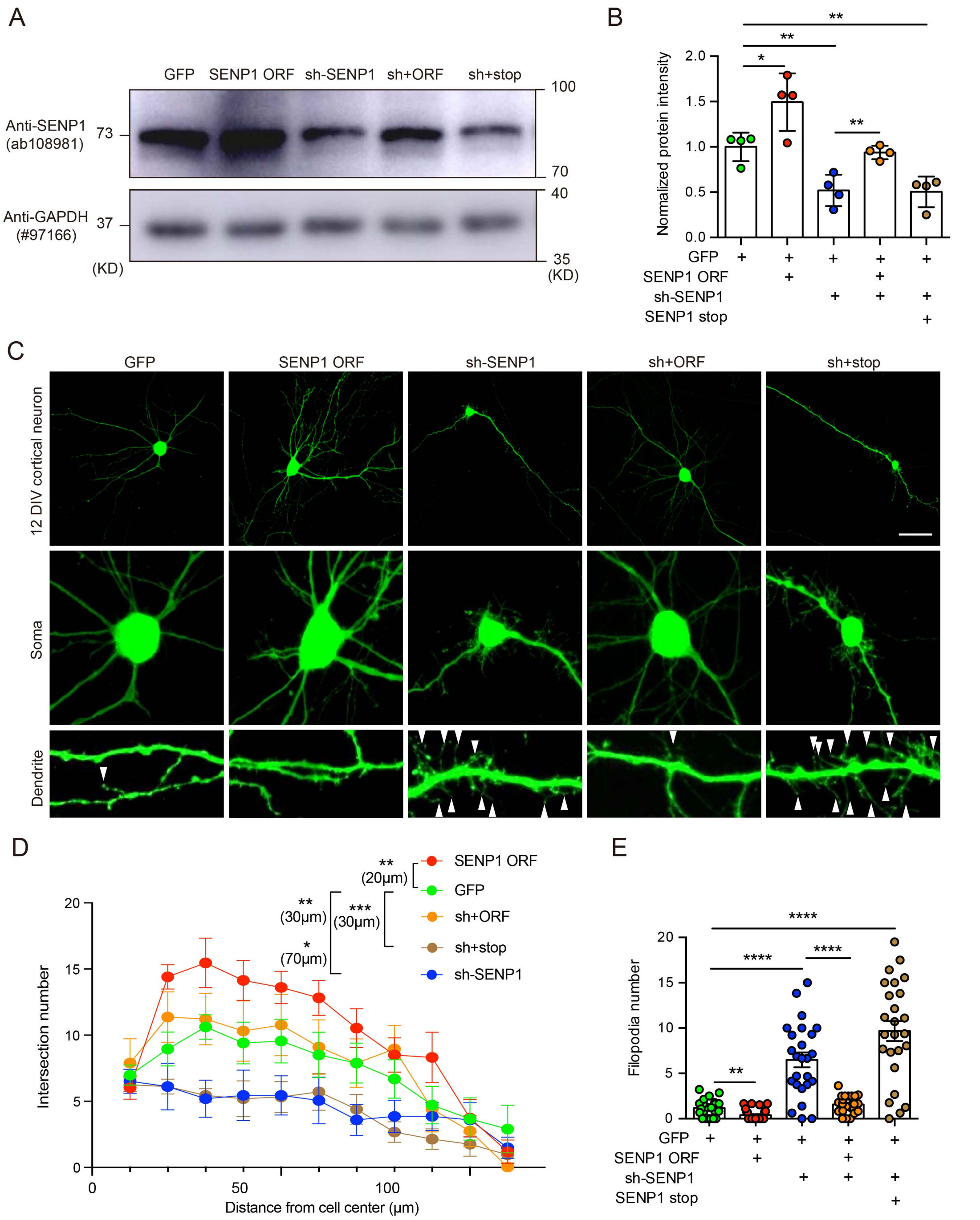


**Figure S4 SENP1 knockdown lead to defects in dendritic morphology and immature spines.**

(A) Representative western blot pictures illustrating protein levels of SENP1 and GAPDH in cultured cortical neurons. Neurons were electroporated with GFP blended with empty vector (GFP), or human SENP1 ORF (SENP1 ORF), mouse SENP1 shRNA (sh-SENP1), mixture of mouse SENP1 shRNA with human SENP1 ORF (sh+ORF), and mixture of mouse SENP1 shRNA with human SENP1 stop-codon-gained ORF (sh+stop), respectively.

(B) Quantification of normalized protein levels for each plasmid combination (from 4 independent batches of cultured cortical neuron).

(C) Representative immunofluorescence staining pictures for cultured cortical neuron at 12 day *in vitro* (for each plasmid combination). The arrowhead indicated the filopodia from dendrite part of neuron. Scale bar, 100μm.

(D) Quantification of Sholl-analysis calculated intersection number of branched dendrites according to distance from cell center for each plasmid combination (n=7, for GFP; n=12, for SENP1 ORF; n=8, for sh-SENP1; n=7, for sh+ORF and n=8, for sh+stop).

(E) Quantification of filopodia number for each plasmid combination (n=23, from 7 neurons for GFP; n=27, from 12 neurons for SENP1 ORF; n=25, from 8 neurons for sh-SENP1; n=23, from 7 neurons for sh+ORF and n=25, from 8 neurons for sh+stop). * $p < 0.05$, ** $p < 0.01$, *** $p < 0.001$, **** $p < 0.0001$. Bars represent means ± SD for Figure S4B; bars represent means ± SEM for S4D and S4E.

Kan Yang et al

Figure S5

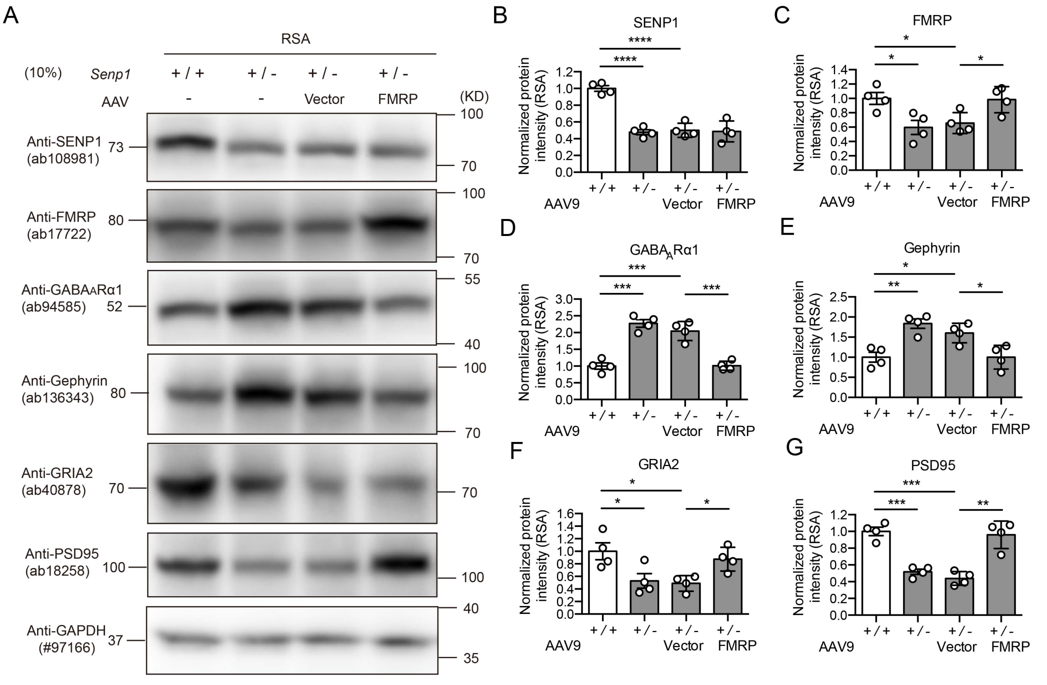


**Figure S5. AAV delivery of FMRP in RSA region rescues inhibitory and excitatory synaptic proteins.**

(A) Representative western blot pictures illustrating protein intensity of SENP1, FMRP, GABA$_A$Rα1, Gephyrin, GRIA2, PSD95 and GAPDH for RSA tissues of mice.

(B) Quantification of normalized protein intensity of SENP1 (n=4, for each genotype).

(C) Quantification of normalized protein intensity of FMRP (n=4, for each genotype).

(D) Quantification of normalized protein intensity of GABA$_A$Rα1 (n=4, for each genotype).

(E) Quantification of normalized protein intensity of Gephyrin (n=4, for each genotype).

(F) Quantification of normalized protein intensity of GRIA2 (n=4, for each genotype).

(G) Quantification of normalized protein intensity of PSD95 (n=4, for each genotype).

* $p < 0.05$, ** $p < 0.01$, *** $p < 0.001$, **** $p < 0.0001$. Bars represent means ± SD.
