## supplemental methods and materials for "SENP1 in the retrosplenial agranular cortex regulates core autistic-like symptoms in mice"

### KEY RESOURCES TABLE

| REAGENT or RESOURCE | SOURCE | IDENTIFIER |
|---|---|---|
| Antibodies | | |
| Rabbit anti-FMRP | Abcam | Cat#ab17722; RRID: AB_2278530 |
| Rouse anti-FMRP | Sigma-Aldrich | Cat#SAB4200597 |
| Mouse anti-GAPDH | Cell Signaling Technology | Cat#97166; RRID: AB_2756824 |
| Chicken anti-GFP | Abcam | Cat#ab13970; RRID: AB_300798 |
| Rabbit anti-PSD95 | Abcam | Cat#ab18258; RRID: AB_444362 |
| Mouse anti-neuroligin 1 | Santa Cruz Biotechnology | Cat#sc-365110; RRID: AB_10708272 |
| Rabbit anti-SENP1 | Abcam | Cat#ab108981; RRID: AB_10862449 |
| Mouse anti-Somatostatin | Santa Cruz Biotechnology | Cat#sc-55565; RRID: AB_831726 |
| Rabbit anti-Parvalbumin | Abcam | Cat#ab181086 |
| Rabbit anti-TBR1 | Abcam | Cat#ab183032 |
| Rabbit anti VGAT | Sigma-Aldrich | Cat#ab2257; RRID: AB_1587623 |
| Chicken anti-Gephyrin | Abcam | Cat#ab136343 |
| Mouse anti-$GABA_AR\alpha1$ | Abcam | Cat#ab94585; RRID: AB_10675844 |
| Guinea pig anti-VGlut1 | Sigma-Aldrich | Cat#ab5905; RRID: AB_2301751 |
| Mouse anti-PSD95 | Thermo | Cat#MA1-046; RRID: AB_2092361 |
| Rabbit anti-GRIA2 | Abcam | Cat#ab40878 |
| Goat anti-GFP | Abcam | Cat#ab5450; RRID: AB_304897 |
| Rabbit anti-GFP | Abcam | Cat#ChIP Grade ab290; RRID: AB_303395 |
| Rabbit anti-PTEN | Cell Signaling Technology | Cat#9552; RRID: AB_10694066 |
| Donkey anti-rabbit, Alexa Fluor 555 | Thermo Fisher Scientific | Cat#A-31572; RRID: AB_162543 |
| Donkey anti-rabbit, Alexa Fluor 488 | Thermo Fisher Scientific | Cat#A-21206; RRID: AB_2535792 |
| Donkey anti-rabbit, Alexa Fluor 647 | Thermo Fisher Scientific | Cat#A-31573; RRID: AB_2536183 |
| Donkey anti-mouse, Alexa Fluor 555 | Thermo Fisher Scientific | Cat#A-31570; RRID: AB_2536180 |
| Donkey anti-mouse, Alexa Fluor 488 | Thermo Fisher Scientific | Cat#A-21202; RRID: AB_141607 |
| Donkey anti-mouse, Alexa Fluor 647 | Thermo Fisher Scientific | Cat#A-31571; RRID: AB_162542 |
| Donkey anti-chicken, Alexa Fluor 488 | Biotium | Cat#20166; RRID:AB_10854387 |

| | | |
|---|---|---|
| Donkey anti-chicken, Alexa Fluor 633 | Biotium | Cat#20168; RRID: AB_10853143 |
| Donkey anti-goat, Alexa Fluor 647 | Thermo Fisher Scientific | Cat#A-21447; RRID: AB_2535864 |
| Donkey anti-guinea pig, Alexa Fluor 555 | Biotium | Cat#20276; RRID: AB_10871473 |
| Donkey anti-mouse, IgG H&L (HRP) | Abcam | Cat#ab6820; RRID: AB_955438 |
| Donkey anti-goat, IgG H&L (HRP) | Abcam | Cat#ab6885; RRID: AB_955423 |
| Donkey anti-rabbit, IgG H&L (HRP) | Abcam | Cat#ab6802; RRID: AB_955445 |
| Bacterial and Virus Strains | | |
| AAV9-CAG-FLEX-EGFP | Taitool Bioscience, Co, Shanghai | N/A |
| AAV9-hSyn-Cre | Taitool Bioscience, Co, Shanghai | N/A |
| AAV9-hSyn-EGFP | Institute of Neuroscience Gene Editing Core | N/A |
| AAV9-hSyn-SENP1-EGFP | Institute of Neuroscience Gene Editing Core | N/A |
| AAV9-hSyn-FMRP-EGFP | Institute of Neuroscience Gene Editing Core | N/A |
| Chemicals, Peptides, and Recombinant Proteins | | |
| DAPI | Thermo Fisher Scientific | Cat#D1306; RRID: AB_2629482 |
| Toloniumchloride | Biossci | Cat#BP034; RRID: N/A |
| OCT | SAKURA | Cat#4583; RRID: N/A |
| Tween-20 | Sangon | Cat#A100777; RRID: N/A |
| Triton-100 X | Sangon | Cat#A110694; RRID: N/A |
| Loading buffer | Applygen | Cat#B1012; RRID: N/A |
| Experimental Models: Organisms/Strains | | |
| Mouse: C57BL/6J | Slac Laboratory Animal | N/A |
| Mouse: *Senp1+/-* | | N/A |
| Software and Algorithms | | |
| ImageJ | NIH | https://imagej.net/Welcome; RRID: SCR_003070 |
| GraphPad Prism | GraphPad Software | https://www.graphpad.com/scientificsoftware/prism/; RRID: SCR_002798 |
| EthoVision XT | Noldus | https://www.noldus.com/ethovision-xt; RRID: SCR_000441 |

### RESOURCE AVAILABILITY

#### Lead Contact

#### Materials Availability

All unique/stable reagents generated in this study are available from the Lead Contact without restriction.

#### Data Availability

All relevant data are available upon request.

### EXPERIMENTAL MODEL AND SUBJECT DETAILS

#### Animals

Male C57BL/6N and *Senp1* +/- mice at the age of 2-4 months old were used for experiments. C57BL/6N mice were purchased from SLAC laboratory (Shanghai). *Senp1* +/- mice were acquired from Jinke Cheng's lab at Shanghai Jiao Tong University. The genotype of *Senp1* +/- mice was determined by performing two parallel PCRs using the same forward primer from exon 8 of *Senp1* (5′-AGTGCACATTCCCTGTCATCTGGT-3′) with different reverse primers: one primed in intron 8 of the *Senp1* gene (5′-TGGGCTGAGTGAGCTTTGACTCTT-3′) and the second primed within the trapping construct (5′-AATCAACTTTGGAGACATGCGGGC-3′). PCR was carried out using standard techniques. All animals were housed under a reversed 12:12 h light-dark cycle with water and food *ad libitum* in the animal facility at the Institute of Neuroscience. All experiments were approved by the Animal Care and Use Committee of the Center for Excellence in Brain Science & Intelligence Technology, Chinese Academy of Sciences, Shanghai, China (IACUC No. NA-016-2016).

### METHOD DETAILS

#### Diagnosis for high-functioning autism

Participants were diagnosed with ASD according to the Diagnostic and Statistical Manual of Mental Disorders, Fifth Edition (DSM-5); confirmed diagnosis with the Autism Diagnostic Interview-Revised (ADIR) and/or Autism Diagnostic Observation Schedule (ADOS). Severity of ASD was defined according to Childhood Autism Rating Scale (CARS): Severe symptoms of ASD were participants whose total score exceeded 36 and who had a rating of 3 or higher on at least 5 of the 15 items of CARS. DD/ID = Developmental delay/intellectual disability: By intellectual assessment, the patients with developmental quotient (DQ) < 75 assessed using Gesell development scales, and intelligence quotient (IQ) < 70 assessed using WISC-R or WPPSI (Wechsler Intelligence Scale for children) were diagnosed as developmental delay or intellectual disability (DD/ID).

#### Behavioral test

Adult male *Senp1* +/+ and *Senp1* +/- mice were tested in the following sequence: open field locomotion, three-chamber sociability test, novel object cognition, restrictive repetitive behavior test and Barnes maze test. Behavioral assays were recorded using cameras. The experimenter handled the mice for constitutive 4 days, 4 min each time to familiarize the mice with the smell of the experimenter before the test. In the fifth day, mice were put into the open field ($40 \times 40$ $cm^2$) for 10 min and the movement of mice was recorded. In the next week, mice were test for three-chamber sociability and novel object cognition. The test apparatus ($40 \times 60$ $cm^2$) contained three adjacent chambers side by side. In the first day mice were tested for social approach. Before the three-chamber experiment started, mice were put into the middle chamber for 10 min to habituate. Then a novel male mouse (stranger A) was put into one of the side chambers for 10 min. The time the subject mouse spent in each chamber was recorded and analyzed. In the next day mice were tested for social novelty. Stranger A and a novel male mouse (stranger B) were put into two side chambers for 10 min respectively, and the time the subject mouse spent in each chamber was recorded and analyzed (Noldus, EthoVisionXT). In novel object cognition test, strange mice were replaced with novel objects. A week later, mice were test for restrictive repetitive behavior. 36 black glass marbles were arranged in a symmetrical 6 × 6 grid on top of 7-cm deep bedding in a clean standard mouse cage (40 × 40 $cm^2$). Mice were put into the cage for 3 min to habituate, then placed in the cage for a 10-min exploration period. The camera took a picture at the beginning and end of the exploration to count the number of buried marbles. 'Buried' was defined as greater than 50% covered by bedding. The next day after marble burying test, mice were placed into a clean standard mouse cage (40 × 40 $cm^2$) for recording cumulative time spent in grooming in the next 30 min. The Barnes maze test was carried out next week. The maze was consisted of a circular platform, 2 m in diameter with 20 evenly spaced holes at the edges. Mice were trained to find a specific small dark chamber under the platform which was the only habitable one called the "escape box." In day 0, mice were placed on the platform for 3 min to habituate followed by pre-training for 3 min to find the escape box and then rest for 1 min. In day 1, mice were placed on the center of the platform for free exploration. The number of errors made to find the escape box was recorded within 3 min followed by 1 min rest. Mice were trained in 2 sessions daily with an inter-trial interval of 1 min for 12 days. The number of errors and latency time made before finding the escape hole were noted. AAV-injection mice were allowed for 1 month to recovery after surgery prior to behavioral test.

**Immunofluorescence assays**

*Senp1 +/+* and *Senp1 +/-* mice were anaesthetized with sodium pentobarbital (Sigma, Cat#P3761, 50 mg/kg) and perfused with 0.1 M PBS followed by 4% PFA. After perfusion, brains were post-fixed overnight in 4% PFA at 4°C and sequentially dehydrated in 15% and 30% sucrose/PBS solution respectively. Then brains were embedded in Optimum Cutting Temperature formulation (OCT) (SAKURA, Cat#4583) and sectioned at 40 μm from parietal lobe to occipital lobe with a Microtome Cryostat (Leica, CM1950) at -25°C. Floating brain sections (40 μm) were rinsed in PBS then blocked overnight at 4°C in PBS containing 5% Bovine albumin (BSA) and 0.2% Triton X-100, followed by incubating with mouse anti-FMRP primary antibodies (Sigma-Aldrich, Cat#SAB4200597, 1:1000) at 4°C for overnight and donkey anti-mouse Alexa Fluor 555 secondary antibodies (Thermo Fisher Scientific, Cat#A-31570, 1:1000) at 4°C for 1 h. All primary and secondary antibodies were diluted with PBS containing 5% BSA and 0.4% Triton X-100. All brain sections were finally counter-stained with DAPI (Sigma, Cat#d9542, 5mg/mL, 1:1000). Sections were washed 3 × 10 min in PBS before incubating with secondary antibodies. For other antibody combinations, sections were rinsed with PBS, blocked and treated with primary and secondary antibodies as described above (see also KEY RESOURCES TABLE). The same immunofluorescence process was applied for cells cultured *in vitro*. Images were captured by objective fluorescent microscope (Olympus, VS120, 10 ×) or confocal microscope (Olympus, FV3000 IX83, 10 ×, 20 × and 60 ×).

**Viruses**

AAV9-CAG-FLEX-EGFP (Serotype 2/9, titer $2\times10^{14}$ vg/mL, vector genome per mL) and AAV9-hSyn-Cre (Serotype 2/9, titer $6.9\times10^{13}$ vg/mL) were purchased from Taitool Bioscience, Co, Shanghai. AAV9-hSyn-EGFP (Serotype 2/9, titer $8.5\times10^{14}$ vg/mL), AAV9-hSyn-SENP1-EGFP (Serotype 2/9, titer $4.1\times10^{13}$ vg/mL) and AAV9-hSyn-FMRP-EGFP (Serotype 2/9, titer $6.2\times10^{13}$ vg/mL) were purchased from Institute of Neuroscience Gene Editing Core.

**Stereotactic injection of AAV virus**

After the anesthetized with sodium pentobarbital (Sigma, Cat#P3761, 50 mg/kg), viruses were injected into the retrosplenial agranular cortex (RSA) bilaterally according to standard mouse brain atlas (Paxinos and Franklin Mouse Brain Atlas, 2nd edition) at the following coordinates: anteroposterior (AP), -1.46 mm; mediolateral (ML), 0.5 mm; dorsoventral (DV), -0.45 mm; anteroposterior (AP), -2.5 mm; mediolateral (ML), 0.75 mm; dorsoventral (DV), -0.5 mm angled 90° toward the midline in the coronal plane. The same standard mouse brain atlas were used for anterior cingulate cortex (ACC): anteroposterior (AP), 0.26 mm; mediolateral (ML), 0.25 mm; dorsoventral (DV), -0.86 mm. Virus stocks were diluted to $1 \times 10^{12}$ vg/ml, and 0.2 ml virus was injected with a speed of 30 nl/min by a micro-injector and micro-infusion pump (PHD 2000, Harvard Apparatus). For the sparse labeling experiment, the AAV9-hSyn-Cre was diluted to $1 \times 10^{8}$ vg/ml, and the AAV9-CAG-FLEX-EGFP was diluted to $1 \times 10^{13}$ vg/ml. About 0.2 ml mixed virus was injected into RSA with a speed of 30 nl/min. After injection, the mice were kept on a warm blanket to maintain the body temperature until fully awake.

**Slice electrophysiological recording**

Mice were anesthetized with sodium pentobarbital (Sigma, Cat#P3761, 50 mg/kg). The cardiac perfusion was performed with -4 °C artificial cerebrospinal fluid (aCSF) (in mM, 125 NaCl, 3 KCl, 2 $CaCl_2$, 2 $MgSO_4$, 1.25 $NaH_2PO_4$, 1.3 $NaH_2PO_4$, 1.3 Na-pyruvate, 26 $NaHCO_3$, and 11 glucose, at pH = 7.4, 290-310 mOsm) saturated with 95% $O_2$ and 5% $CO_2$. The mouse brain was removed rapidly and softly, then submerged into cold aCSF for ~ 1 min. Coronal slices (300 μm) were dissected and incubated in a chamber containing aCSF at room temperature for ~ 1 h before recording. Slice was transferred into the recording chamber which was constantly perfused with aCSF, the temperature was controlled at ~ 30 °C by the temperature controller (Warner instrument cooperation, USA). Whole-cell recordings were performed on pyramidal neurons in the layer II/III of RSA, the neurons were visualized by infrared microscope (Andor) equipped with epifluorescence and infrared-differential interference contrast (DIC) illumination. Patch pipettes were pulled from borosilicate glass (3-5 MΩ) and filled with a pipette solution consisting of (in mM), 130 K-gluconate, 20 KCl, 10 HEPES, 0.2 EGTA, 4 $Mg_2$ATP, 0.3 $Na_2$GTP, and 10 $Na_2$-phosphocreatine, at pH 7.3 (290-310 mOsm). During the recording of mEPSCs and mIPSCs, 1μm TTX was added into the circulating aCSF. While measuring mIPSCs, the membrane potential was hold at 0 mV to minimize the mEPSCs. While recording mEPSCs, the membrane potential was hold at -70 mV to minimize the mIPSCs. Data were acquired with pClamp9.2 (Molecular Devices) using an AxonMultiClamp 700A amplifier (Molecular Devices), low pass filtered at 2 kHz, and digitized at 20-100 kHz (Digidata 1322A; Molecular Devices). Before recording, the junction potential was corrected. The data analysis was performed with the Clamfit 10.3. All chemicals were purchased from Sigma.

**Western blot and co-IP assays**

To assess protein intensity *in vitro*, neurons from wild-type brains were dissected at E14.5 for primary cell culture and electroporation. To assess protein intensity *in vivo*, the embryonic cortex and RSA of 3-month mice were dissected. After being homogenized at 1000 rpm for 1 min, tissues were lysed by ultrasound for 30 min in ice-cold RIPA lysis buffer. Following a further ultrasound lysis for 30 min, the samples were centrifuged at 12000 rpm for 10 min at 4 °C. For Western blot, proteins in supernatant were combined with loading buffer (Applygen, Cat#B1012) and was heated to 100 °C for 30 min. The protein lysate was electrophoresed using 8%-12% SDS-PAGE gels and ran for 100 min at 120V. The proteins were transferred to a transfer membrane (Millipore Immobilon-P, Cat#IPVH00010) for 100 min at 200 mA. The membrane was blocked with 5% TBST containing Bovine albumin (BSA), washed 3×10 min with 1×TBST with 0.1% Tween-20 (Sangon, Cat#A100777), and then was incubated for 12 h with the primary antibody at 4 °C. The membrane was washed 3×10 min with 1×TBST with 0.1% Tween-20, incubated with the secondary antibody for 1 h. Signals were detected using an Ai600 system following manufacturer's protocol. For co-Immunoprecipitation, 650 μl supernatants were added with 20 μl untreated and pre-blocked protein G-agarose beads for 1 h upside down at 4 °C. Proteins with beads were centrifuged at 12000 rpm for 10 min at 4 °C, and supernatants were incubated with primary antibodies or their corresponding IgGs as IP control for 4 h upside down at 4 °C and then for 1h at 4 °C with 30 μl of pre-blocked protein G-agarose beads. Following 12000 rpm for 10 min at 4 °C, precipitates attached by remaining 20 μl beads were added with 40 μl loading buffer, and proteins were eventually eluted by boiling the beads to 100°C for 30 min before SDS-PAGE.

**Primary neuron culture and transfection**

Cortical tissues of E14.5 C57BL/6 mice were dissected, followed by mincing and digestion with 0.125% trypsin (Thermo Fisher, Cat#15090046) to generate dissociated neurons for cell culture. Neurons were treated either under electroporation in the suspension status with the AMAXA Nucleofector (AAD-1001S, Germany) in 6-well plates, or under lipofection with Lipofectamine-2000 in 12-well plates. For electroporation, AMAXA Mouse Neuron Nucleofector kits, together with Nucleofector Solution buffer were applied for resuspension of the neurons. Program of "mouse, neuron, 0-005" on the AMAXA nucleofector II device was feasible, with $3\times10^6$ cells to 3 mg of plasmid for each electroporation as an optimal condition. For Lipofectamine-2000-mediated transfections, a considerably small quantity of both plasmid and Lipofectamine-2000 (0.1 mg plasmid with 0.1 ml Lipofectamine-2000 for $2\times10^5$ cells in one well of the 12-well-plate) was appropriate for avoiding toxic effects of Lipofectamine-2000. After being fed with Neurobasal medium supplemented with 2% B27 (Life Technology, Cat#17504044), cells were fixed at 12 DIV for immunofluorescence staining, or harvested at 12 DIV for Western blot.

**Transmission electron microscopy assay**

In order to obtain RSA tissue samples, a 3-month mouse was perfused with 1% glutaraldehyde and 4% paraformaldehyde in PBS. The RSA was dissected and treated with the method mentioned in Figure 6B. Tissue samples were cut into 1-$mm^3$ size for further treatment with 1% osmic acid in PBS and gradient dehydration with alcohol. After being treated with a mixture of epoxypropane (Sinopharm Chemical Reagent, Cat#80059118) and resin (Electron Microscopy Sciences, Cat#14900), samples were embedded with pure resin, and then sliced by a diamond tool bit. Target cells were randomly selected and captured with the transmission electron microscope (Nippon Tekno, JEOL-1230, Japan).

### QUANTIFICATION AND STATISTICAL ANALYSIS

**Statistical Analysis**

Data were plotted as mean ± standard error of mean (SEM) or Standard Deviation (SD). Statistical tests were analyzed with GraphPad Prism (GraphPad Software). For comparisons between two groups, data sets were analyzed with Student's t-test (two-tailed, paired or unpaired). For comparisons among data of more than two groups in Figure 2F, one-way ANOVA was used. *$p < 0.05$, **$p < 0.01$, ***$p < 0.001$, ****$p < 0.001$.
